# Characterisation of organised smooth endoplasmic reticulum suggests a route towards synthetic compartmentalisation

**DOI:** 10.1101/2022.10.27.514093

**Authors:** Andras Sandor, Marketa Samalova, Federica Brandizzi, Verena Kriechbaumer, Ian Moore, Mark D Fricker, Lee J Sweetlove

## Abstract

Engineering of subcellular compartmentalisation is one of synthetic biology’s key challenges. Among different approaches, *de novo* construction of a synthetic compartment is the most coveted but also most difficult option. Restructuring the endoplasmic reticulum (ER), via the introduction of recombinant oligomerising ER-membrane resident proteins, is an alternative starting point for building a new compartment. The presence of such proteins leads to a massive expansion of the ER and the formation of organised smooth endoplasmic reticulum (OSER), a large membranous compartment. However, OSER is poorly characterised and our understanding of its effect on the underlying biology of the plant is limited. Here we characterise a range of OSER compartments and show how the structure of the inducing polyprotein constructs affect the final compartment morphology, with the cytosolic-facing antiparallel oligomerisation domain demonstrated to be an essential component to trigger OSER formation. We show that while OSER retains a connection to the ER, a diffusional barrier exists to both the ER and the cytosol. Using high-resolution quantitative image analysis, we also show that the presence of this large compartment does not disrupt the rest of the ER network. Moreover, transgenic *Arabidopsis* constitutively expressing the compartment-forming polyproteins grew and developed normally. These properties collectively suggest that OSER could be developed as a plant synthetic biology tool for compartmentalisation, combining the benefits of several existing strategies. Only a single protein construct is necessary to induce its formation, and the compartment retains a delimiting membrane and a diffusional barrier to the rest of the cell.

## Introduction

Subcellular compartmentalisation is a fundamental principle of life on Earth, allowing the spatiotemporal separation of distinct cellular functions from the environment and other compartments (Bolsover *et al*., 2011; Monnard and Walde, 2015). The membranous organelles of eukaryotes are the typical examples of such structures (e.g., the nucleus, mitochondria or the endoplasmic reticulum (ER)), but membraneless organelles are also broadly used by both eukaryotes and prokaryotes for a variety of purposes: for example, protein-shell based bacterial microcompartments are widespread structures to contain the catabolism of toxic compounds (Agapakis, Boyle and Silver, 2012; Kerfeld *et al*., 2018).

Compartmentalisation allows different metabolic pathways to be physically separated and quasi-independently regulated by modulating substrate availability and the subcellular microenvironment (Boeynaems *et al*., 2018). This allows mutually incompatible pathways (e.g., due to different solution requirements, competition for substrates or futile cycling) to exist in the same cell (Sweetlove and Fernie, 2013), and permits compartments to specialise in certain metabolic processes (Sweetlove and Fernie, 2018). Furthermore, toxic intermediates or final products generated by specialised cellular metabolism can be sequestered into compartments, protecting the bulk of the cell (Sweetlove and Fernie, 2013; Cárdenas, Almeida and Bak, 2019).

Due to these benefits, the engineering of *de novo* synthetic compartments has long been an attractive target for synthetic biology. While repurposing pre-existing organelles is considered simpler (e.g., by re-targeting enzymes to them utilising specific signals) and has produced several successes in improving the production of compounds of interest in plants (e.g., dhurrin (Henriques de Jesus *et al*., 2017), artemisinin (Malhotra *et al*., 2016), terpenes (van Herpen *et al*., 2010; Liu *et al*., 2011; Eljounaidi *et al*., 2014)), this approach can face challenges from competing endogenous pathways in their new environment, which were optimised by millions of years of evolution (Polka, Hays and Silver, 2016; Gao and Zhou, 2019). Synthetic new compartments can reduce this issue. For example, liquid-liquid phase separated membraneless organelles have recently become a popular alternative approach (Alberti, 2017; Uversky, 2019). Their popularity is fuelled by the limited number of constructs required to engineer them with often a single scaffold molecule sufficient to trigger the formation of membraneless organelles (Nott *et al*., 2015; Bracha, Walls and Brangwynne, 2019), and specific cargo-capture (Banani *et al*., 2016; Reinkemeier, Girona and Lemke, 2019; Garabedian *et al*., 2021) and optogenetic control methods now exist to improve their utilisation (Bracha, Walls and Brangwynne, 2019; Zhao *et al*., 2019). However, while the lack of a delimiting membrane eliminates some engineering challenges (e.g., the requirement for specific pores or channels), it also reduces control over the local microenvironment and permeability (Polka, Hays and Silver, 2016; Schuster *et al*., 2018).

A completely synthetic membranous compartment could in theory be superior to the aforementioned compartmentalisation strategies. Due to its lack of native functions, a synthetic membranous compartment would have limited crosstalk with the host background, and the presence of a membrane would facilitate control over the contents of the compartment. However, this approach requires a much higher level of understanding of the biology of the host, making it a far more difficult strategy compared to re-purposing existing compartments or utilising membraneless organelles. While some progress has been reported in recent years in this field (e.g., by the use of the maize storage protein γ-Zein to build functionalised vesicles (Reifenrath *et al*., 2020)), there is still no broadly usable method for the generation of synthetic compartments for metabolic engineers.

One potential candidate to serve as the basis for developing a new compartment is the ER. This is the starting point for the membrane endogenesis pathway involved in the formation of several organelles (Stefano and Brandizzi, 2018), and itself is highly dynamic and undergoes large-scale restructuring (Almsherqi *et al*., 2009; Sparkes, Hawes and Frigerio, 2011; Stefano and Brandizzi, 2018). ER-membrane resident proteins (such as the reticulons (Voeltz *et al*., 2006; Shibata *et al*., 2008), atlastins (Chen *et al*., 2011; Ueda *et al*., 2016) and Lunapark proteins (Kriechbaumer *et al*., 2018)) influence the architecture of the ER, facilitating interconversion between different ER morphologies (tubules, three-way junctions and cisternae, respectively).

The ability of the ER to change shape allows it to take on some highly specialised forms. One of these is the appearance of tightly stacked arrays of smooth ER sheets, forming concentrated whorls or regular sinusoidal arrays with cubic symmetry, collectively known as organised smooth endoplasmic reticulum (OSER) (Snapp *et al*., 2003; Borgese, Francolini and Snapp, 2006; Sandor *et al*., 2021). While OSERs were primarily investigated in yeast and mammalian cell lines, studies utilising recombinant fluorescent proteins to explore the secretory pathway of plant cells have revealed much about how OSER is formed; for a detailed review on this topic see Sandor *et al*. (2021). These discoveries were mostly serendipitous, as oligomerising membrane-bound proteins often accidentally trigger the formation of OSER (Yamamoto, Masaki and Tashiro, 1996; Snapp *et al*., 2003; Barbante *et al*., 2008). As such, much that was uncovered about OSER was incidental and studies were not done in a systematic way since OSER was often not the main focus of the researchers (Sandor *et al*., 2021). To employ OSER as potential compartmentalisation tool in plant cells, an in-depth characterization of this structure and knowledge on the molecular underpinnings needed to modulate its formation seem necessary.

Here, we report an in-depth characterisation of OSER structures formed from a range of ER-membrane-resident protein constructs to assess the relationship between the constitutive domains of the OSER-inducing protein and the OSER structure whose formation it triggers. We show that the OSER compartment has a diffusional barrier to both the ER and the cytosol. Quantitative analysis of the dynamics and morphology of the remaining ER shows that it is not significantly disrupted by the formation of the OSER structure. These findings, together with the capacity to constitutively or transiently express OSER in plants without any obvious negative phenotype suggest that OSER could be utilised as a compartmentalisation tool in plant cells and would be a valuable asset to the growing field of plant synthetic biology.

## Results

### Serendipitous discovery of an ER-remodelling polyprotein

To interrogate trafficking in the plant endomembrane system, a fluorescent polyprotein probe was developed, consisting of a 22 amino acid transmembrane domain (BP22 – derived from the BP80 vacuolar sorting receptor from *Pisum sativum* (Paris *et al*., 1997; Brandizzi *et al*., 2002)) flanked by a N-terminal GFP and a C-terminal YFP (due to this organisation, the construct was named G22Y) targeted to the ER (Figure 1A). While the construct was expected to pass through the endomembrane system to the plasma membrane as it was previously reported when GFP was genetically fused to the BP22 domain (Brandizzi *et al*., 2002), when transiently expressed in mature tobacco leaves, this double-tagged construct instead integrated into the ER forming large (up to 25 μm in diameter), self-organising compartments as apparent from the GFP and YFP signal observed by fluorescent confocal microscopy (Figure 1B). Notably, these G22Y structures showed a clear helical patterning (Figure 1B). To assess the formation of the G22Y structures, a time-course experiment was carried out imaging the structures every 12 h until 168 h after transient expression (Supplementary Figure S1). The G22Y signal consistently started to appear around 36 h as small (up to 5 μm) compartments which aggregated into a single (or, in rare cases, two) large compartment per cell by 60 h. Cells with compartments remained viable up to 6 weeks, demonstrating the stability of these compartments.

**Figure 1:**
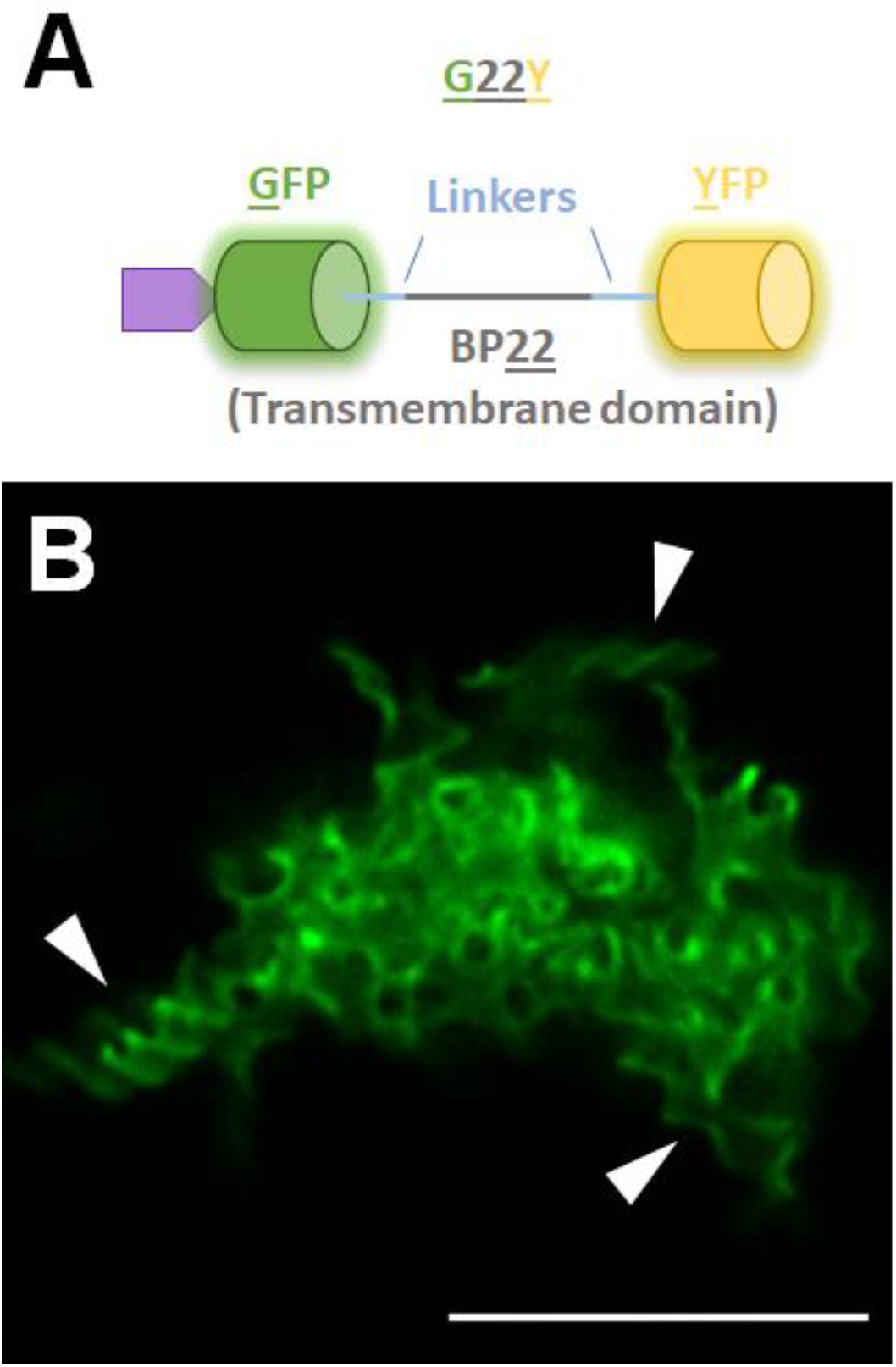
Remodelling the ER using the synthetic oligomerising polyprotein G22Y. A: Schematic of the G22Y polyprotein, composed of two dimerising fluorescent proteins (GFP and YFP) flanking a transmembrane domain (BP22) and a N-terminal ER targeting signal peptide. B: Fluorescence confocal microscopy images of the structures formed in mature *N. tabacum* leaves 7 days after agroinfiltration. Arrowheads point to some notable helical organisations in the structure. The image was captured using high-resolution Airyscan imaging of the structures. Scale bar: 10 μm.

To confirm that the compartments are derived from the ER, they were co-expressed with RFP-HDEL (Nelson, Cai and Nebenführ, 2007) and TAR2-RFP (Kriechbaumer, Botchway and Hawes, 2016), two fluorescent markers for the ER lumen and ER membrane, respectively. The overlapping signals suggested that both the lumen and the membrane of the compartment are contiguous with the ER (Figure 2A). To determine if the compartment retains connection to the ER network, fluorescence recovery after photobleaching (FRAP) was used to assess lateral diffusion between the ER and the G22Y compartment, by photobleaching compartments co-expressed with RFP-HDEL or the bulk ER (control). The ER lumenal marker rapidly recovered into the compartment, confirming that it was still connected to the ER. However, the rate of recovery of fluorescence was significantly reduced when compared to the rest of the ER, suggesting a diffusional barrier between the bulk of the ER and the lumen of the G22Y compartment (Figure 2B).

**Figure 2:**
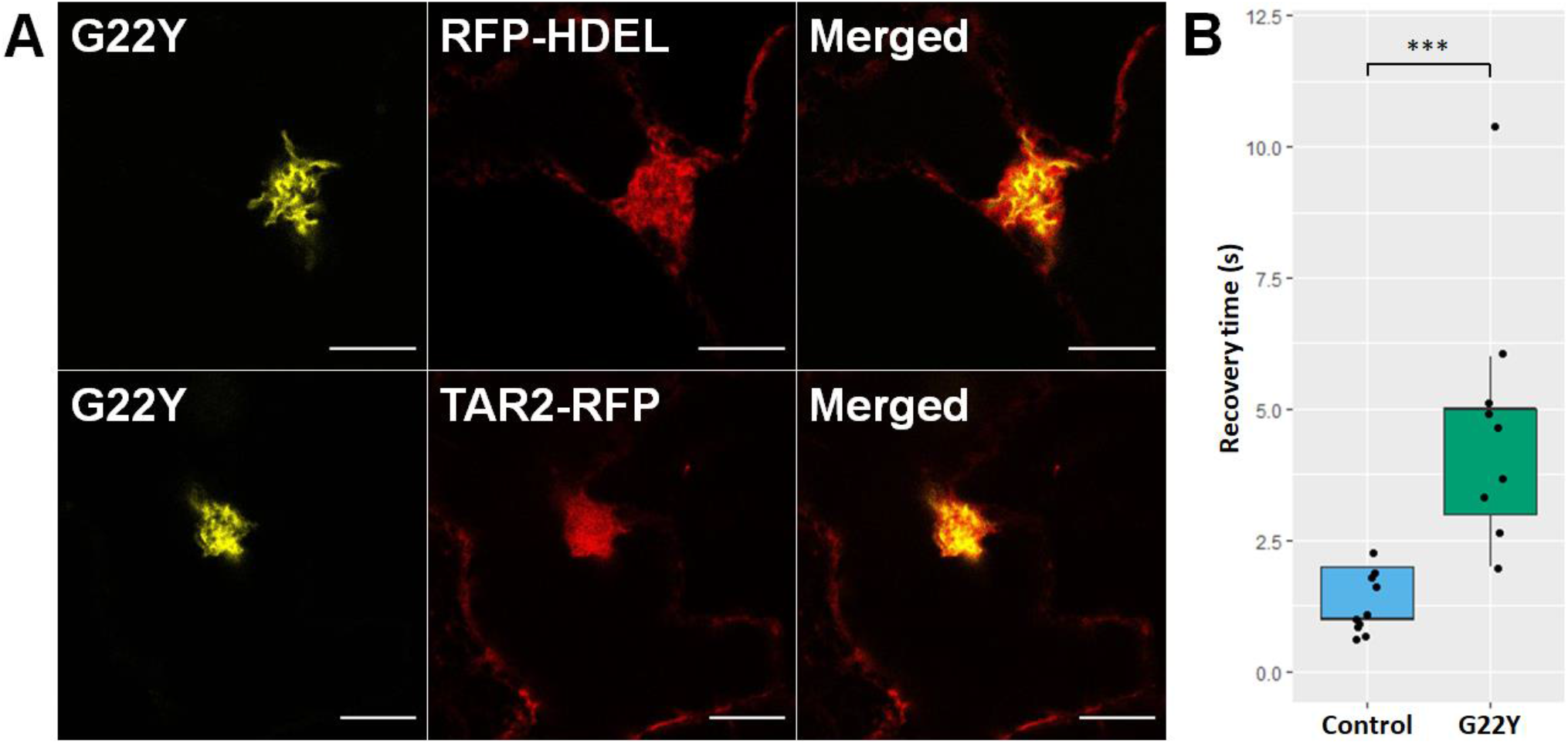
Co-expression of G22Y with fluorescent ER markers shows the compartment’s connection to the ER membrane and lumen. A: G22Y (yellow) was co-expressed with the fluorescent ER-luminal marker protein RFP-HDEL and ER-membrane marker TAR2-RFP (both in red). The signals overlap, confirming that the G22Y compartment derives its membrane and lumen from the ER. The fluorescence confocal microscopy images were captured 7 days after agroinfiltrating mature *N. tabacum* leaves. Scale bars: 10 μm. B: The recovery rate of RFP-HDEL after FRAP of the G22Y compartment is slower when compared to the periperhal ER (Control). Asterisks depict significance levels: *** for p < 0.001, n=9.

### Expanding the suite of compartment-forming polyproteins to investigate the structural basis of the OSER compartment

The formation of the G22Y compartment with its unusual helical organisation and diffusional barrier to the lumen of the ER warranted further exploration. To determine how the protein domains affected the formation of the compartment, a range of five alternative G22Y-derived constructs were generated.

First, to assess the requirement of a cytosolic or ER-lumenal facing dimerising domain for the formation of the compartment, we removed either dimerising fluorescent proteins, which yielded the G22 and 22Y constructs, (Figure 3). In two further constructs, the removed fluorescent protein was replaced with a synthetic dimerising coiled-coil domain CC-Di (Fletcher *et al*., 2012), which is of similar length to GFP or YFP, but has a much stronger binding affinity (K_D_ < 10^−8^ M) (Fletcher *et al*., 2012) and dimerises in parallel, in contrast to GFP and YFP, which dimerise at lower affinity (K_D_ ≈ 10^−4^ M) in an antiparallel fashion (Zacharias *et al*., 2002). These two constructs (C22Y and G22C) enabled us to explore the effects of different binding orientations and strengths of the scaffold proteins on the compartment. The final construct G-Y(cyt) had its transmembrane domain and ER-targeting signal peptide removed to determine if ER membrane integration is essential for the formation of this compartment (Figure 3).

**Figure 3:**
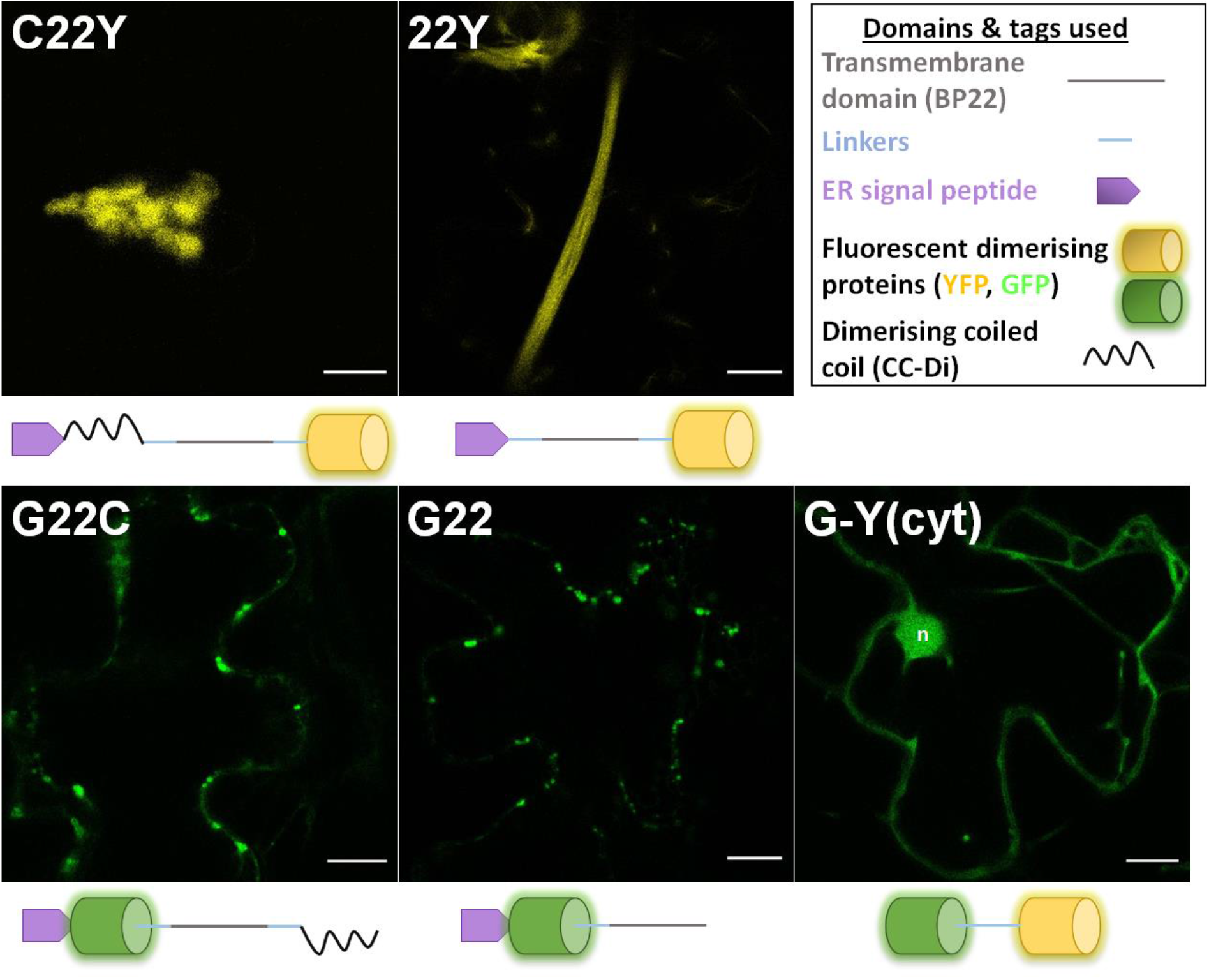
Visualising alternative G22Y-derived constructs to expand the compartment-forming scaffold toolkit. Five new polyproteins were designed, with two having either the N-terminal YFP or the C-terminal GFP removed (G22 and 22Y, respectively). In two constructs, the removed fluorescent dimerising protein was replaced with a synthetic dimerising coiled-coil domain (G22C and C22Y) and in one construct the transmembrane domain and the ER-targeting signal peptide was removed (G-Y(cyt)). Top right box shows the constitutive domains of the schematics of G22Y-derived constructs. Confocal microscopy images of the new constructs 7 days after agroinfiltrating mature *N. tabacum* leaves. Yellow: YFP. Green: GFP. n: nucleus. Scale bars: 10 μm.

Fluorescent confocal microscopy imaging of transiently transformed tobacco leaves revealed that two of the constructs formed compartments of similar (or in some cases larger) size to the previously described G22Y compartments: 22Y often presented more elongated structures, while C22Y showed more globular compartments. G22 and G22C fluorescent signal accumulated in numerous smaller (< 5 μm) compartments. G-Y(cyt) did not induce the formation of any compartments and appeared to be located mainly in the cytosol and nucleus (Figure 3). Cells with compartments showed normal cell behaviour (e.g., cytoplasmic streaming and ER network rearrangements identical to untransformed cells (Supplementary Video SV1)) until leaf senescence, highlighting the unlikely toxicity of the compartments.

To further determine the subcellular localisation of the fluorescent signal, the new constructs were co-expressed with a range of fluorescent markers. Co-expression of C22Y and 22Y with the ER-lumenal RFP-HDEL and ER-membrane-localised TAR2-RFP revealed co-localisation of the two fluorescent signals, confirming that the compartment’s membrane is derived from the ER. The cytosolic marker Peredox-mCherry (Hung and Yellen, 2014) also partially co-localised with the C22Y and 22Y compartments, suggesting the presence of trapped cytosol in or around the compartment structure (Supplementary Figures S2, S3). These markers also highlighted the formation of several spherical vacuoles in close proximity to 22Y compartments (Supplementary Figure S4).

Co-expression of G22C and G22 with RFP-HDEL revealed no overlapping signal between the small compartments and the ER lumen, and the co-expression of G22C with the peroxisomal marker YFP-PEX2 revealed that the small compartments are likely peroxisomes, suggesting a mis-localisation of the polyproteins instead of forming an ER-derived compartment (Supplementary Figure S5).

### Further investigation into the C22Y and 22Y structures

The two variants construct forming large ER-derived compartments, C22Y and 22Y, were further investigated using high-resolution airy-scanning confocal microscopy and transmission electron microscopy (TEM) to observe the detailed structures of the compartments (Figure 4).

**Figure 4:**
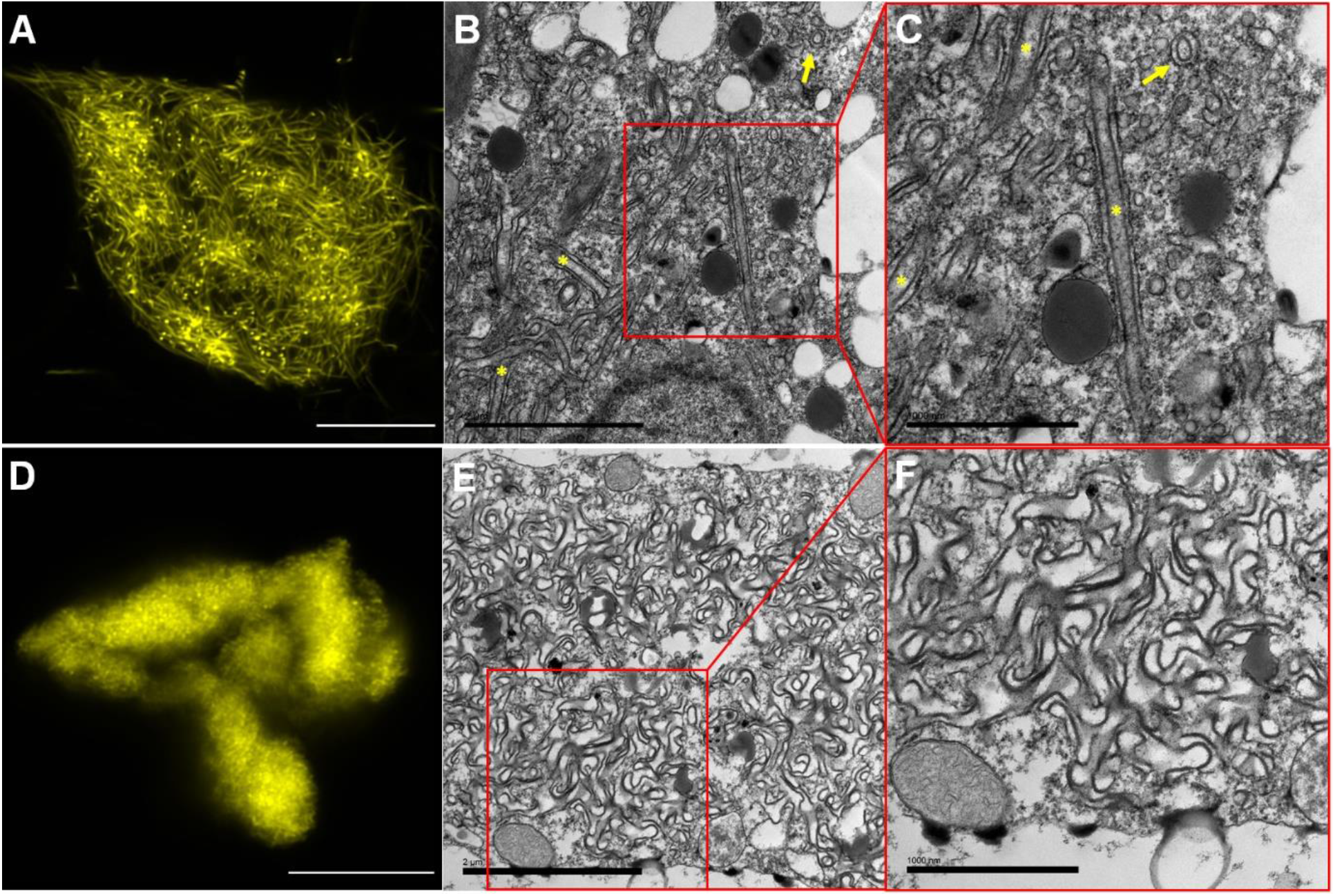
High-resolution characterisation of the 22Y and C22Y compartment structures. A & D: High-resolution Airyscan fluorescent confocal microscopy images of 22Y (A) and C22Y (D) compartments. Images were captured 7 days after agroinfiltration of mature *N. tabacum* leaves. Scale bars: 10 μm. B, C, E and F: TEM images of 22Y (B and C) and C22Y (E and F) compartments. 22Y structures often form two parallel membrane pairs (yellow asterisks). This is likely an ER cylinder, with some cross-sections of these highlighted with a yellow arrow. Scale bars: B & E: 2 μm, C & F: 1 μm.

The two constructs induced the formation of morphologically distinct compartments. 22Y compartments were composed of several long fibrous strands, likely double-membraned ER cylinders with cytosol trapped in the centre (Figure 4A-C). C22Y compartments showed a more homogenous fluorescent signal, which could not be further resolved using Airyscan confocal microscopy (Figure 4D). TEM images of C22Y compartments presented complex patterns of internal organisation, with densely packed membrane structures (Figure 4D-F), which are likely too dense for clear resolution using confocal imaging.

To determine if the two new compartment types retain their connection to the ER, and if they have a similar diffusional barrier as the G22Y compartments (Figure 2), FRAP was carried out by co-expressing the compartment-forming proteins with RFP-HDEL. When photobleached, the RFP-HDEL signal recovered rapidly, but significantly slower than for the peripheral ER in cells not expressing OSER constructs (control), suggesting that both the 22Y and C22Y compartments are connected to the ER, but a diffusional barrier exists (Figure 5A).

**Figure 5:**
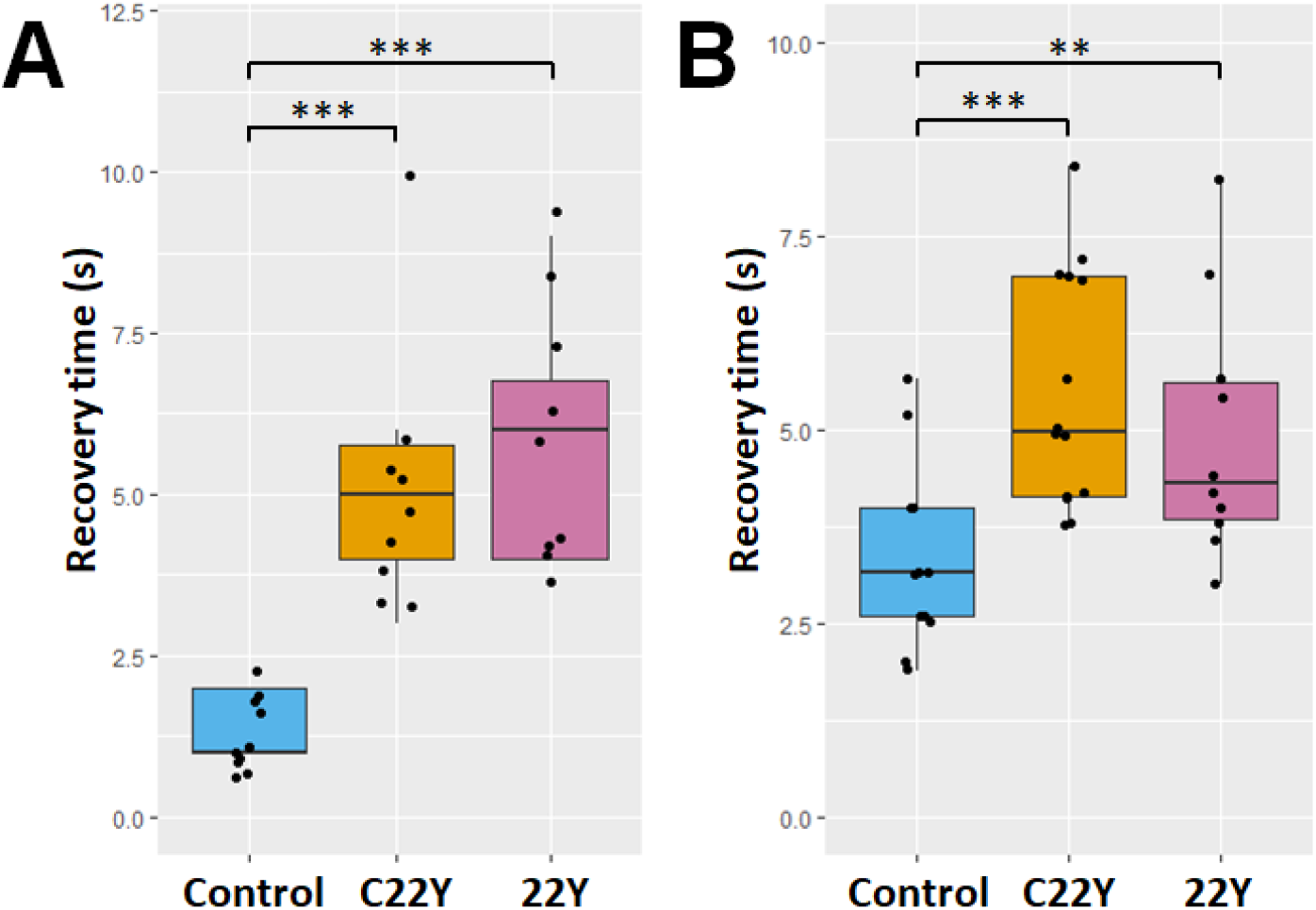
Lumenal and cytosolic fluorescence recovery after photobleaching rates are reduced in C22Y and 22Y compartments. FRAP experiments on the C22Y and 22Y compartment types using ER lumenal RFP-HDEL (A) and cytosolic Peredox-mCherry (B) both show a significantly increased recovery time when compared to the respective controls, suggesting a diffusional barrier between the compartment and both the ER lumen and the bulk cytosol. Asterisks depict significance levels: ** for p < 0.01 and *** for p < 0.001. n = 9 for A and n = 13 for B for all constructs. The boxes represent the interquartile range, the horizontal line in the box shows the median, and the whiskers the minimum and maximum values (excluding outliers).

As co-expression with Peredox-mCherry suggested that these compartments have cytosol trapped within their complex membrane structures (Supplementary Figure S2, S3), a FRAP experiment using Peredox-mCherry was also carried out. This showed similar results to the RFP-HDEL FRAP, suggesting a diffusional barrier to exist between the bulk cytosol and the cytosol trapped by the compartments (Figure 5B).

The presence of the diffusional barrier to the rest of the cell means that molecules can be effectively compartmentalised temporarily by these structures. However, to determine if the compartment-forming scaffold proteins are also sequestered primarily in the compartment, the proportion of the total cellular fluorescence inside the compartments were measured for C22Y. This showed that the scaffold proteins almost exclusively reside in the compartment (an average of 92.7% of total cellular fluorescence was present in the compartment – Supplementary Figure S6).

### Quantitative analysis of the architecture and dynamics of the ER

As the G22Y, C22Y and 22Y compartments are derived from the ER, their substantial size raised the possibility of a significant disruption to the host cell’s ER. To determine if this was the case, the remaining ER was visualised in cells containing one of the three compartments using co-infiltrated RFP-HDEL. Initial visual inspection showed that non-compartmentalised ER retains a normal phenotype. However, a more accurate method for determining the potential ER disruption was needed. To this end, short (20 s) time-series were captured using high-resolution Airyscan confocal microscopy focusing only on the peripheral ER (the compartments were not imaged) and quantitatively analysed using the AnalyzER software package (Pain *et al*., 2019). 41 parameters were measured and analysed including topological, morphological and kinetic parameters of the ER cisternae, tubules and the polygonal regions enclosed by them (Table S1). ANOVAs of the 41 parameters showed only a single one of the parameters (the average persistency of cisternae) was significantly different between the WT and any of the compartment types (Table S1), but this significance disappeared when accounting for multiple testing. This suggests that the formation of the OSER compartments did not significantly disrupt the structure and dynamics of the rest of the ER in the compartment-carrying cells.

### Expressing the compartment in stably transformed *A. thaliana*

To determine if the compartments can be developed and maintained in stably transformed *A. thaliana* as well as with transient expression, the two constructs which produced the largest compartments, C22Y and 22Y were introduced under the drive of the strong constitutive cauliflower mosaic virus 35S promoter (Odell, Nagy and Chua, 1985). T1 seedlings were selected based on antibiotic resistance and inspected for fluorescence, and two independent T2 lines were isolated for each construct (Figure 6). Seedlings showed YFP fluorescence in all tissue types, but T2 seedlings had noticeably stronger signal in their roots. Transformed plants developed similarly to WT controls and set seed normally.

**Figure 6:**
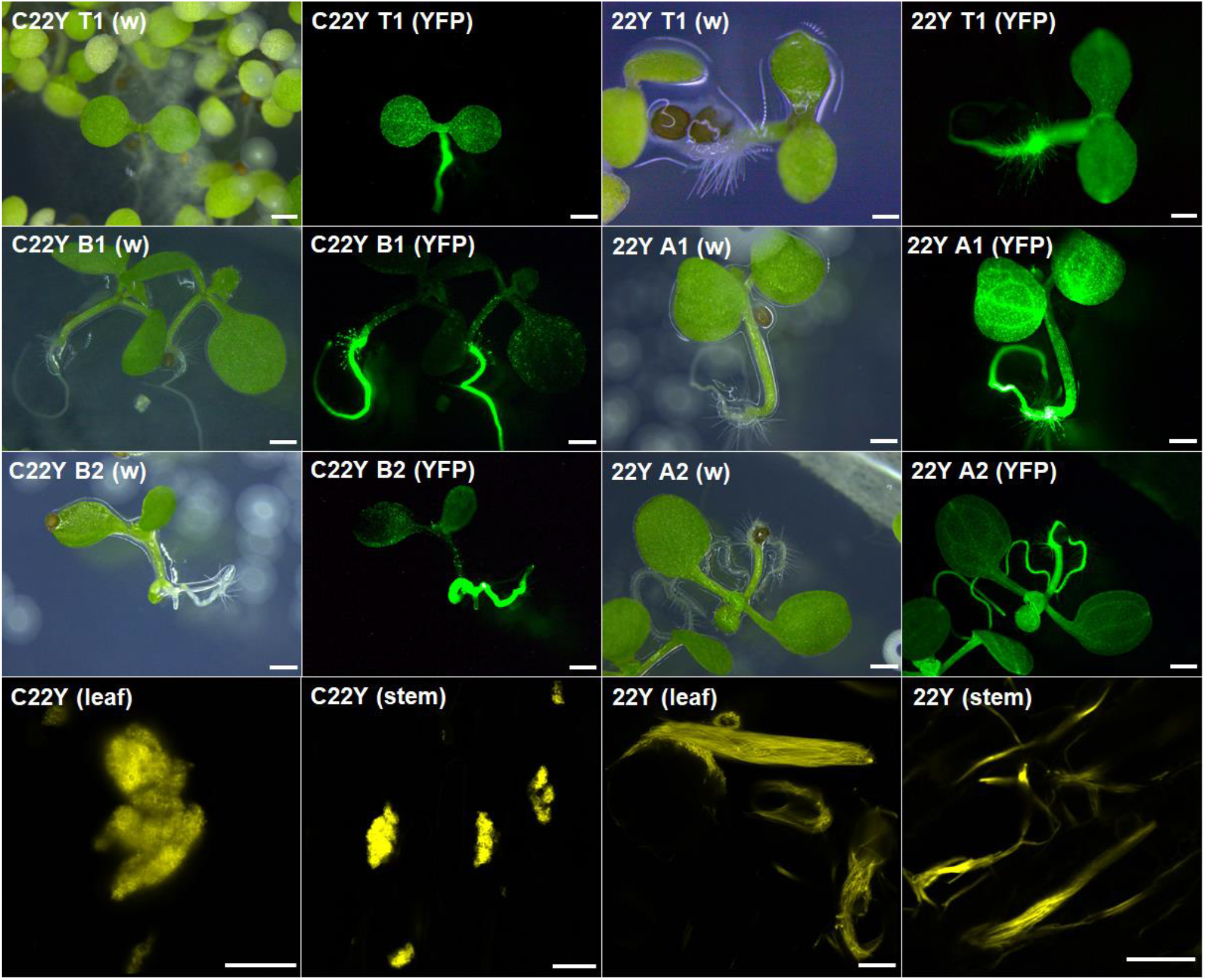
Stably transformed *A. thaliana* lines form C22Y and 22Y compartment structures similar to those obtained by transient expression in tobacco. Top three rows: fluorescent whole seedling imaging of 5-6 d old stably transformed *A. thaliana* on MS-media plates. Pairs of images show the white-light (w) and YFP fluorescence (YFP) of T1 (top row) and T2 (second and third row) plants. Fluorescent signal is present in all tissues but is noticeably stronger in the roots of T2 plants. Scale bars: 1 cm. Bottom row: high-resolution Airyscan laser confocal microscopy images of C22Y and 22Y expressing 6 weeks old T2 plants. Both compartment types show identical morphology to the previously described compartments following transient expression of each construct. Scale bars: 10 μm.

Six-week-old T2 *A. thaliana* plants were imaged using fluorescence confocal microscopy to determine if compartments formed and if their morphologies were identical to what was observed in transiently transformed tobacco epidermis. The stably transformed plants showed fluorescence in all cells in the stem and leaves, and fluorescent signal was detected in seeds as well (Supplementary Figure S7). High-resolution Airyscan confocal microscopy images confirmed in the stem and leaves of stably transformed *A. thaliana* plants that the C22Y and 22Y compartments have a phenotype consistent with the transiently expressed compartments described above (Figure 6).

## Discussion

Building subcellular compartments is one of the central challenges of synthetic biology due to the numerous applications that require separation of functions from the bulk of the cell. Here we have shown that OSER, which was previously primarily viewed as a disruption of ER morphology, can be used as a viable tool for this purpose in plants. The structures described above had a diffusional barrier to the bulk of the cell and the lumen of the ER, providing effective compartmentalisation, without significantly disrupting the rest of the ER.

### Effects of the dimerising domain topology on compartment structure

From our experiments it was clear that the changes made to the dimerising domains used in the synthetic polyprotein constructs had a great impact on the observed compartment morphology.

In our study, a cytosol-facing antiparallel dimerising domain was essential, as removal or replacement of it with a parallelly binding domain led to mislocalisation to the peroxisomes, instead of the formation of large ER-derived compartments.

The ER lumenal dimerising domain also had a clear effect on the compartment morphology, since its modification (from GFP to YFP or complete removal) lead to two compartments with structures completely different from the G22Y compartments. It is possible that the binding topology (antiparallel, parallel or missing) has an effect on what membranes can be bridged by these polyprotein oligomers, leading to these differences. An antiparall (head-to-head) binding domain is expected to be more flexible, capable of bridging different membranes together in trans, while a parall binding domain is spatially more restricted, likely to only binding to the same membrane in cis. Further studies using more compartment-forming polyproteins could be used to gain more insight into the exact relationship between the oligomerising domain topologies and the induced compartment structures.

### A putative OSER subclass

Similarities between the synthetic compartments and previously described OSER structures are notable enough to warrant comparison. Both classes are derived from the ER, forming large membranous structures which retain some connection to the ER – indeed previous FRAP experiments on OSER also showed a reduced diffusion between the ER lumen and the OSER structure (Marti *et al*., 2010). Furthermore, the formation of OSER did not affect the normal functioning of the ER (Ferrero *et al*., 2015), and the quantitative analysis of the ER network in our study (Table S1) also showed no significant disturbance to the ER. Plants stably transformed with the compartments developed normally, similar to what was reported by previous studies for OSER (Ferrero *et al*., 2015).

Morphologically, the C22Y compartments are also similar to some of the larger OSER structures (Ferrero *et al*., 2015; Sandor *et al*., 2021), and the fibrous 22Y structures resemble elongated lamellae, another typical OSER phenotype (Sandor *et al*., 2021). Notably, 22Y compartments are morphologically identical to the OSER structures described induced by the *nuc* mutation of the MVP1 protein in *A. thaliana* (Jancowski *et al*., 2014). However, the MVP1 is predicted to be a non-membrane resident monomeric protein indirectly involved in ER-to-Golgi export, and the mutation was hypothesised to induce OSER by blocking this export pathway, inadvertently causing the accumulation of other OSER-inducing proteins in the ER (Jancowski *et al*., 2014). By contrast, in this study the results from the quantitative ER analysis using AnalyzER showed no major disruptions to the ER, making it unlikely that 22Y causes the formation of the compartment described in this study in the same manner.

Equally, there are also clear morphological differences between our compartments when compared to OSER: for example, the helical patterning of G22Y compartments have not been described previously in connection with OSER to our knowledge. OSER also usually presents as multiple structures in a cell, unlike in our study, where a single compartment was formed almost exclusively in each cell.

The induction of compartment formation also presents some similarities to OSER formation. OSER can be induced by two types of proteins: ER-resident membrane proteins and proteins disrupting ER-to-Golgi export (Sandor *et al*., 2021). For the former group, there is some controversy in the literature to the necessity of a cytosol-facing oligomerising domain, with some studies showing it to be essential (Yamamoto, Masaki and Tashiro, 1996; Snapp *et al*., 2003; Barbante *et al*., 2008), while others refuting this claim (Marti *et al*., 2010; Jancowski *et al*., 2014; Ferrero *et al*., 2015; Grados-Torrez *et al*., 2021). Recently these opposing views were attempted to be reconciled into a single unified theory of OSER formation, which hypothesised multiple parallel pathways to induce OSER (Sandor *et al*., 2021). In this study, only ER-resident membrane proteins were capable of inducing the formation of our compartment. Furthermore, a cytosol-facing antiparallelly dimerising domain was essential for the formation of the compartments.

Overall, this compartment system seems to have significant similarity to previously described OSER structures in terms of formation and effects on the host cell, but there are some morphological differences as well, suggesting that the structures described above might constitute a subgroup of OSER.

### Potential applications of the synthetic compartment system

The compartment system described in our study has a number of valuable properties that make it a good candidate for functionalisation as a plant synthetic biology tool. No detrimental effects were observed on transiently or stably transformed plants, and the ER of the transformed cells were quantitatively shown to be not significantly disrupted. The compartments show a consistent morphology across two different species, and the constitutive polyprotein scaffold almost exclusively accumulates inside the compartment structure. These properties together suggest that this compartment system could be used to improve the production of recombinant proteins, especially difficult-to-express membrane proteins or toxic proteins which would be mostly sequestered from the bulk of the cell. Proteins of interest could be genetically fused directly to the polyprotein scaffold, or via the introduction of a modular attachment method (e.g., the SpyCatcher – SpyTag system (Reddington and Howarth, 2015) to covalently bind the target recombinant protein to the scaffold before or after release to the cytosol) to potentially allow greater expression of valuable proteins before purification.

Another potential application is to develop this system as a synthetic microdomain. This is enabled by the high specificity of the scaffold molecules to the compartment in combination with the diffusional barrier between the compartment lumen and the ER lumen, as well as the trapped cytosol and the bulk cytosol. Together, these properties are expected to be sufficient to generate enable probabilistic metabolic channelling, where if two subsequent enzymes of a pathway are co-localised at high concentrations (by binding them to the compartment), the intermediate of the pathway is more likely to encounter the next enzyme of the pathway, rather than to diffuse to the bulk cytosol (Sweetlove and Fernie, 2018). This would be a valuable tool for the metabolic engineering of plants, by providing a way to channel flux down a pathway of interest, especially if the intermediate if unstable, toxic or secreted (Sweetlove and Fernie, 2013; Cárdenas, Almeida and Bak, 2019).

A new modular compartmentalisation system would be of great interest for plant synthetic biology, and an OSER-based membranous compartment has numerous advantages. Compared to current popular approaches (co-opting existing organelles, building novel compartments and membraneless organelles) OSER has a number of attractive properties: unlike membraneless organelles, it is delimited by a membrane bilayer allowing greater control over the internal composition and is induced by a single polyprotein construct, making is similarly simple to use. While the compartment retains some connections to the ER, there is a clear diffusional barrier making it a more specific tool than simply targeting proteins to the ER, retaining some of the benefits of a novel compartment and the host ER is not significantly impacted. Overall, these properties give this strategy notable potential as a tool for plant synthetic biology.

## Methods

### Plant husbandry and materials

Ethanol-sterilised *A. thaliana* seeds were grown own MS-agar (4.33 g L^-1^ Murashige & Skoog medium (Duchefa Biochemie, Haarlem, Netherlands), 15 g L^-1^ agar) medium plates with 50 μg mL^-1^ kanamycin and 200 μg mL^-1^ cefotaxime in a growth cabinet with a 16 light - 8 h dark photoperiod (at light intensity of 120 μM m^-2^ s^-1^) at 20 °C. Seedlings were transplanted to soil (3:1 ratio of Sinclair Pro modular seed growing compost (Sinclair Pro, Cheshire, UK) and Sinclair pro fine vermiculite (Sinclair Pro, Cheshire, UK) with 0.4 g L^-1^ Exemptor (ICL, Ipswich, UK) as an insecticide) after 7 days and grown in a greenhouse with a 16 h light – 8 h dark photoperiod at 25 °C, with natural light supplemented to achieve light intensity of up to 200 μM m^-2^ s^-1^ using ATTIS-7 LED grow lights (Plessey, London, UK). *N. tabacum* seeds were sown directly onto soil and grown in a greenhouse in identical condition as above.

### Plant transformation

Transient transformation were carried out as described in Sparkes *et al*. (2006) by pelleting *A. tumefaciens* culture after overnight growth, washing twice using a modified infiltration buffer (5mM MES pH 5.6, 5 mM MgCl_2_, 500 μM acetosyringone), and injecting *A. tumefaciens* suspended (at OD_600_ = 0.05) in infiltration buffer into the underside of 4-6 weeks old *N. tabacum* leaves using a needleless syringe. When multiple protein constructs were co-expressed, every *A. tumefaciens* carrying a different construct was introduced at equal concentrations to the infiltration buffer.

Stable transgenic lines were obtained through transformation by floral dip followed by antibiotic selection (Clough and Bent, 1998).

### Genetic construct design and molecular cloning

Prior to gene synthesis, the protein constructs were modelled using pyMOL (v2.3.4, Schrödinger LLC, New York, USA) to determine the binding orientation of the oligomerising domains. The genetic constructs coding for the polyprotein scaffolds C22Y and G22C were synthesised by Twist Bioscience (San Francisco, USA) and supplied recombined in the Gateway entry vector pTWIST-ENTR. 22Y and G22 were derived from these constructs, respectively, by removing the coiled-coil domain CC-Di using polymerase chain reaction (PCR) via Phusion High-Fidelity DNA Polymerase (Thermo Fisher Scientific, Waltham, USA) according to the manufacturer’s instructions. G-Y(cyt) was generated similarly from G22Y by removing BP22 and the ER-targeting peptide. G22Y was designed and cloned as described in (Samalova, Fricker and Moore, 2006). Briefly, a sp-mGFP5-BP22 product of overlapping PCR was digested with *Spe*I/*Xho*I and inserted into *Spe*I/*Sal*I sites of pVKHEn6-YFP_myc_ vector. The gene parts used for the design of the constructs are described in detail is Supplementary Table S2. The primers used for PCR can be found in Supplementary Table S3.

Constructs in the Gateway entry plasmid pTWIST-ENTR were subcloned into the plant Gateway expression plasmid pK7WG2 (Karimi, Inzé and Depicker, 2002) using the Gateway LR Clonase II Enzyme Mix (Thermo Fisher Scientific, Waltham, USA) according to the manufacturer’s instructions. pK7WG2 plasmids were then introduced into OneShot Mach1 T1 Phage-Resistant Chemically Competent Cells (Thermo Fisher Scientific, Waltham, USA) *Escherichia coli* according to the manufacturer’s instructions. Plasmids were then subsequently transformed into *A. tumefaciens* LBA4404 competent cells using the freeze-thaw method as described in (Wise, Liu and Binns, 2006).

### Confocal laser scanning microscopy

Leaf epidermal samples of *A. thaliana* and *N. tabacum* and seeds and stem samples of *A. thaliana* were imaged using a Zeiss LSM880 confocal microscope equipped with an Airyscan detector (Carl Zeiss AG, Oberkochen, Germany). Images were commonly captured with a C-Apochromat 40x/1.2 W autocorrect M27 water-immersion objective (Carl Zeiss AG, Oberkochen, Germany) at 1024×1024 px resolution at pixel spacing of 20-100 nm with 4-line averaging and 16-bit depth at maximum speed, with typical excitation at 488 nm (GFP), 514 nm (YFP and chloroplast) or 561 nm (RFP and mCherry) and emission at 490-510 nm (GFP), 520-560 nm (YFP), 590-640 nm (RFP and mCherry) or 660-700 nm (chloroplasts). When imaging two fluorophores in one sample, care was taken to avoid spectral overlap by using sequential excitation and line switching.

For experiments focusing on quantitative analysis of the ER network and dynamics and fluorescence recovery after photobleaching (FRAP), the delay between the images in time-series was minimised by reducing image resolution to 512×512 px and using 2-line averaging to give a frame rate of approximately 125 ms image^-1^. When determining what proportion of the scaffold protein constructs were localised to the synthetic ER compartment, whole cell Z-stack images were captured with extra care to avoid any signal saturation. For FRAP experiments, leaf epidermal samples were treated with 25 μM Latrunculin B (Sigma-Aldrich, St. Louis, USA) for 10 min at room temperature to immobilise the ER immediately prior to imaging. Image analysis was carried out in ImageJ (v1.52k) and statistical analysis in R (v3.6.1). Statistical comparison of different groups for FRAP was analysed using Student’s t-test.

RFP-HDEL (Shockey *et al*., 2006), TAR2-RFP (Kriechbaumer, Botchway and Hawes, 2016), Peredox-mCherry (Hung and Yellen, 2014), GFP-TIP1:1 (Nelson, Cai and Nebenführ, 2007) and YFP-PEX2 (Sparkes, Hawes and Baker, 2005) were used as fluorescent probes to visualise the ER lumen, ER membrane, cytosol, tonoplast and peroxisomes, respectively.

### Quantitative analysis of ER network architecture and dynamics

Short time-series Airyscan images (20 sec) were collected of the ER network at the apical end of cells expressing a compartment-forming protein construct and RFP-HDEL. Cells were only imaged if the compartment was not visible, to visualise only the unmodified ER in the region-of-interest. Images underwent quality control in ImageJ (v1.52k) to remove time-series that had drifted in the Z-axis or were blurry. Images drifting in the X or Y axes were corrected (if possible) using the StackReg (v1.0) module (Thévenaz, Ruttimann and Unser, 1998).

The images were then processed and analysed using the AnalyzER software package (v1.1), following the protocols described in Pain *et al*. (2019). The parameters investigated are described in Supplementary Table S1. These parameters were then statistically analysed using MANOVA to correct for multiple comparisons followed by analysis of variance (ANOVA) to determine if there were statistically discriminating features (kinetic or morphological, in the tubules, cisternae or enclosed areas) between the ER of different compartment-expressing cells and cells only expressing RFP-HDEL.

### Transmission electron microscopy

Mature *N. tabacum* leaf sections were fixed, stained using zinc-iodine-osmium and embedded in resin (as described in (Kittelmann, 2018). 90 nm sections were cut from the embedded samples, placed onto 200 mesh copper grids and then post-stained for 5 min with lead-citrate. The stained sections were imaged using a FEI Tecnai T12 transmission electron microscope (FEI, Hillsboro, USA), operated at 120 kW. Images were captured by a GATAN OneView digital camera (AMETEK, Pleasanton, USA) and analysed in ImageJ (v1.52k).

## Acknowledgements

A.S. acknowledges funding from the University of Oxford, the EPSRC & BBSRC Centre for Doctoral Training in Synthetic Biology (grant EP/L016494/1).

We would like to thank Clare Jones (Department of Biological and Medical Sciences, Oxford Brookes University, Oxford, UK) for her help with leaf section staining and embedding and Dr. Errin Johnson (Sir William Dunn School of Pathology, University of Oxford, Oxford, UK) for her help with TEM sectioning and imaging. Finally, we would like to recognise the massive contribution made by the late Ian Moore to this research. The original discovery of the ER-derived compartment was made in his laboratory and it was Ian who championed the idea of developing the compartment for synthetic biology. He was keen for others to take this work on and we would like to think that he would have been pleased with the outcome of the research presented in this manuscript.

## Author contributions

AS designed, carried out and analysed the experiments and wrote the paper. LJS and IM conceived the study. MS designed and carried out the initial experiments to build the G22Y construct and co-wrote the paper. Transgenic Arabidopsis plants were generated at the MSU-DOE Plant Research Laboratory, Michigan State University, under the supervision of FB. LJS, MDF, VK and FB supervised the study and co-wrote the paper

## Supplementary Materials

**Supplementary Figure S1 (below):**
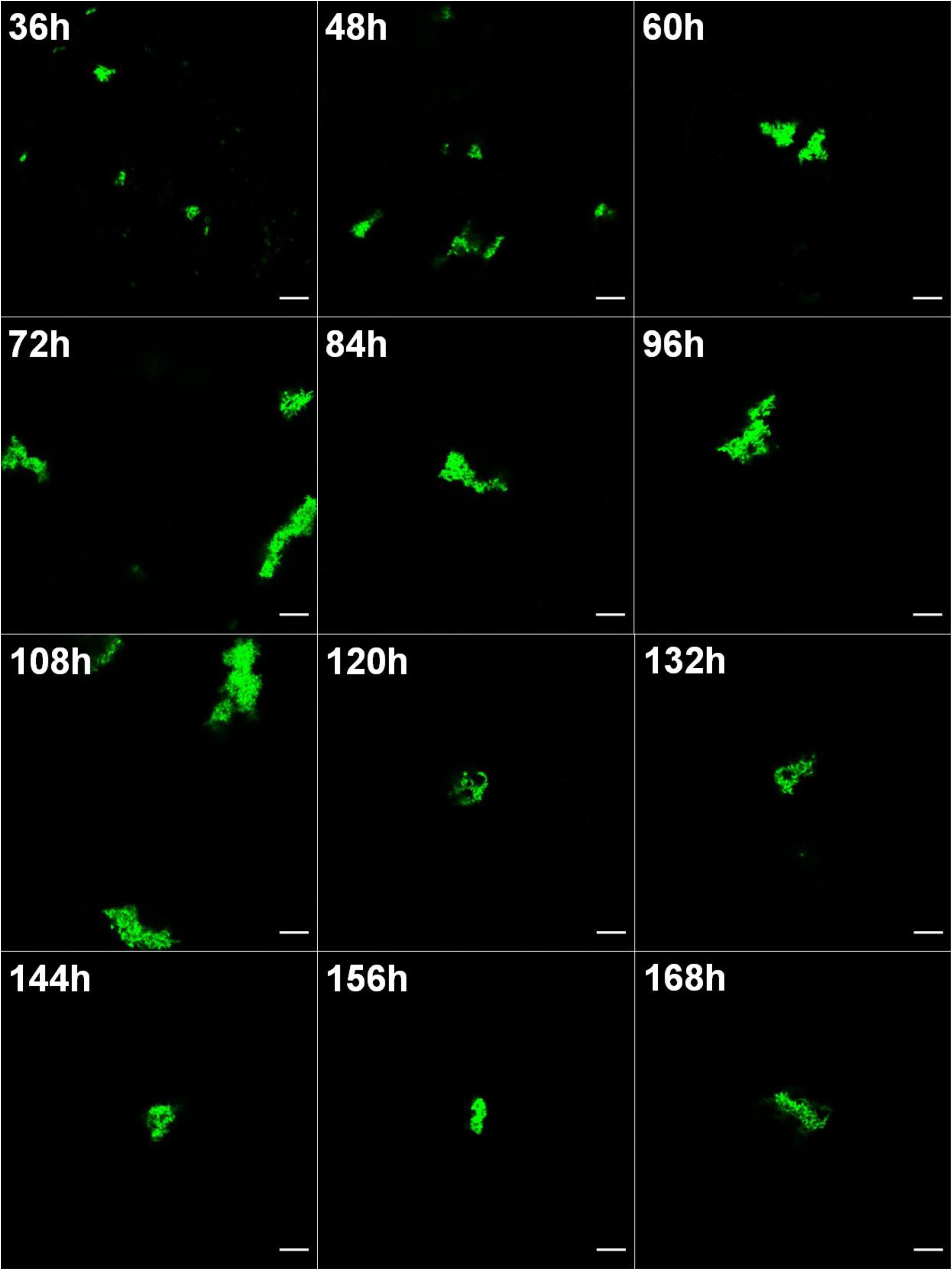
Expression time-course of the G22Y compartments. Fluorescent confocal microscopy images of transiently transformed mature *N. tabacum* leaves shows how several smaller compartments aggregate into one or two large compartments per cell around 48-60 h after agroinfiltration. No expression of G22Y was detected before 24 h. Images shown here were collected from the same plant over the course of a week, and is representative of the morphologies and progression of appearance of compartments. In some cases, fluorescence in compartments was seen up to 6 weeks. Scale bars: 10 μm.

**Supplementary Figure S2:**
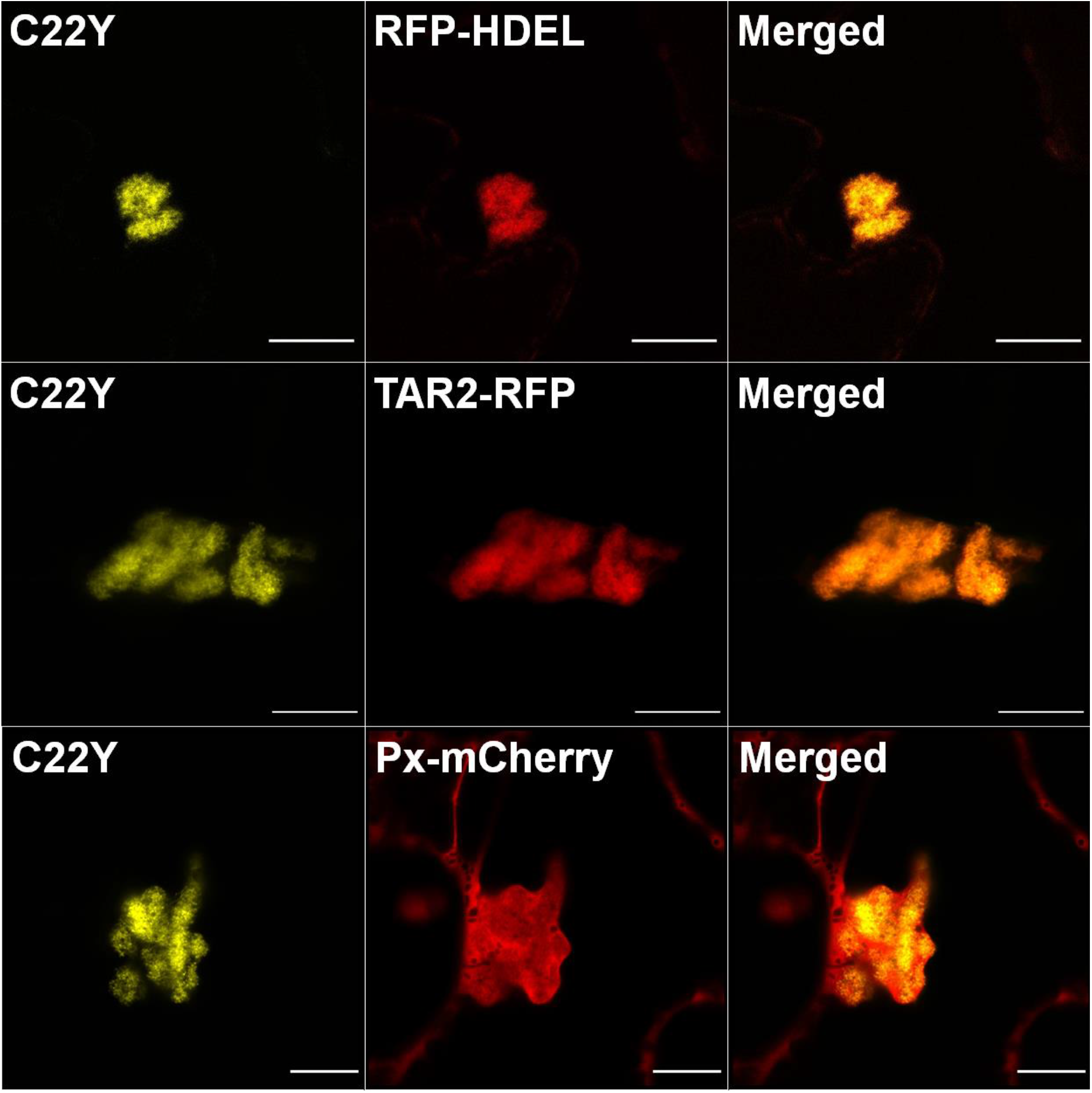
Co-expression of fluorescent markers with C22Y. The three florescent markers RFP-HDEL (ER lumen), TAR2-RFP (ER membrane) and Peredox-mCherry (cytosol) all show co-localisation with the C22Y signal. Images were captured 7 days after agroinfiltration of mature *N. tabacum* leaves. Scale bars: 10 μm.

**Supplementary Figure S3:**
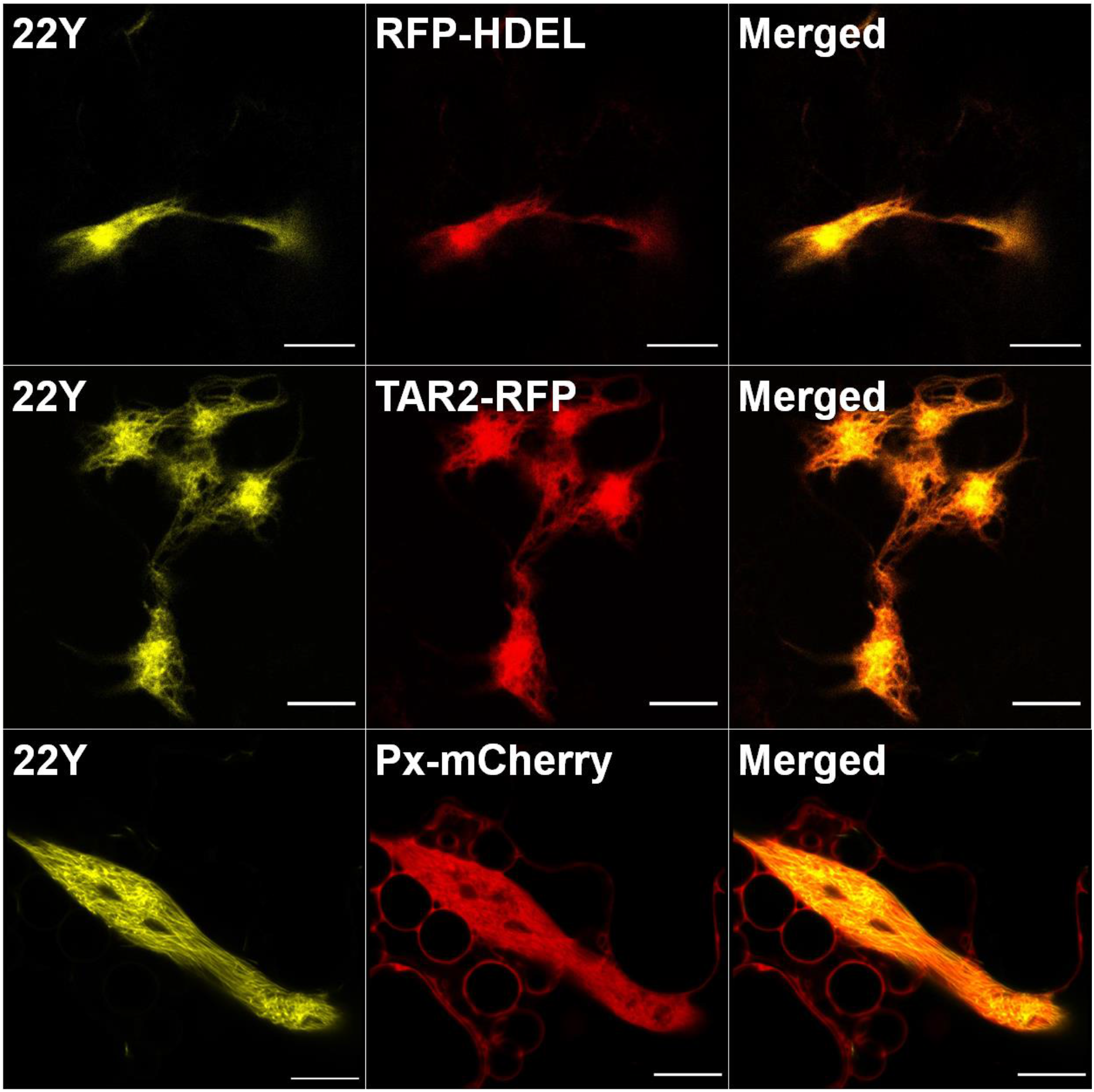
Co-expression of fluorescent markers with 22Y. The three florescent markers RFP-HDEL (ER lumen), TAR2-RFP (ER membrane) and Peredox-mCherry (cytosol) all show co-localisation with the 22Y signal. Images were captured 7 days after agroinfiltration of mature *N. tabacum* leaves. Scale bars: 10 μm.

**Supplementary Figure S4:**
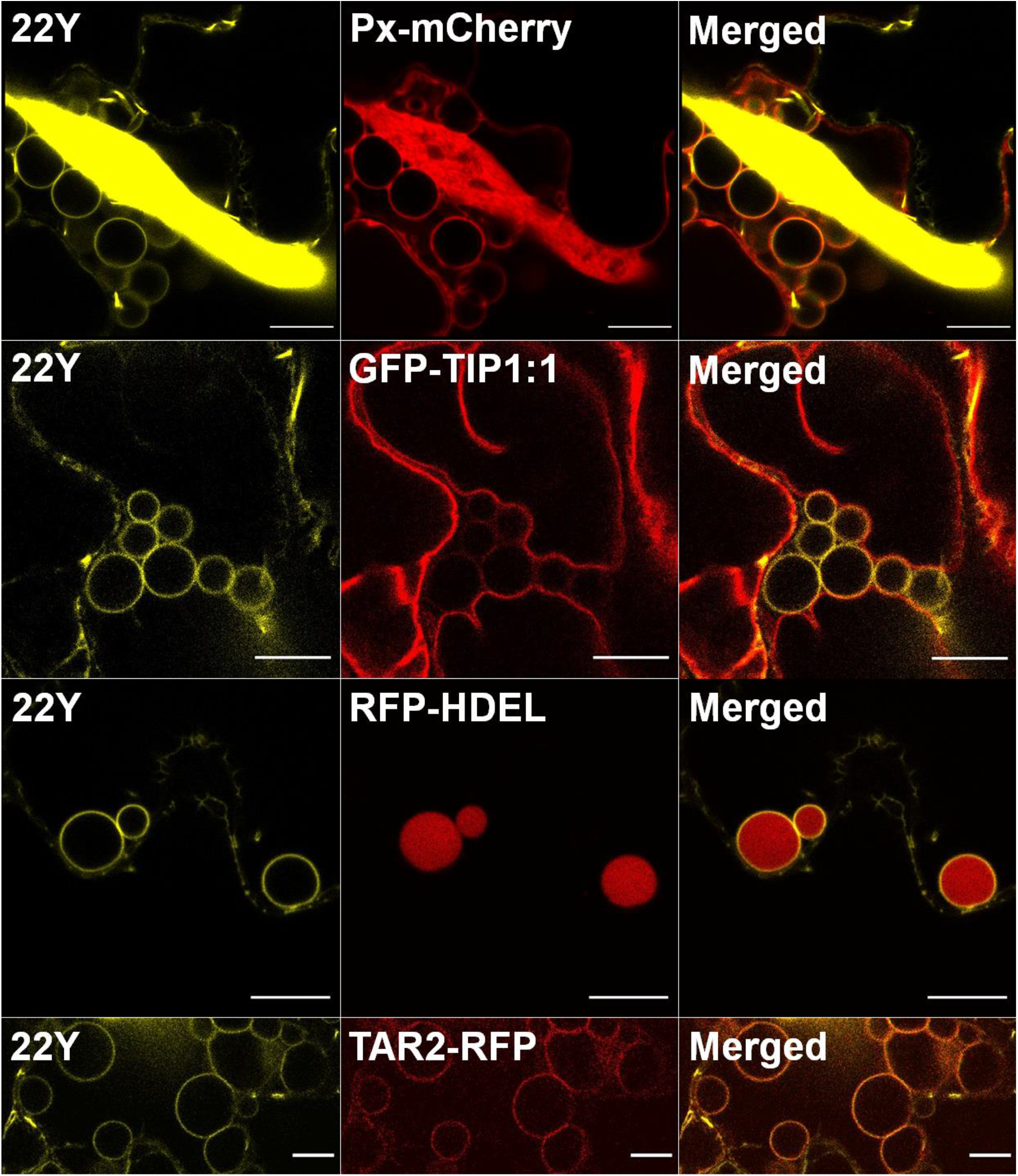
Characterisation of the 22Y spheres. Four fluorescent markers were co-expressed with 22Y and visualised 7 days after agroinfiltration of mature *N. tabacum* leaves. The cytosolic Peredox-mCherry image is a high-contrast version of the one from Supplementary Figure S3, showcasing the presence of trapped cytosol next to the 22Y spheres. The tonoplast membrane marker GFP-TIP1:1 is present in the membrane of the spheres, but at lower concentration than in the tonoplast membrane. The luminal ER marker RFP-HDEL is present in the lumen of the spheres, and the ER membrane marker TAR2-RFP is also present in the membrane of the 22Y spheres. Scale bars: 10 μm.

**Supplementary Figure S5:**
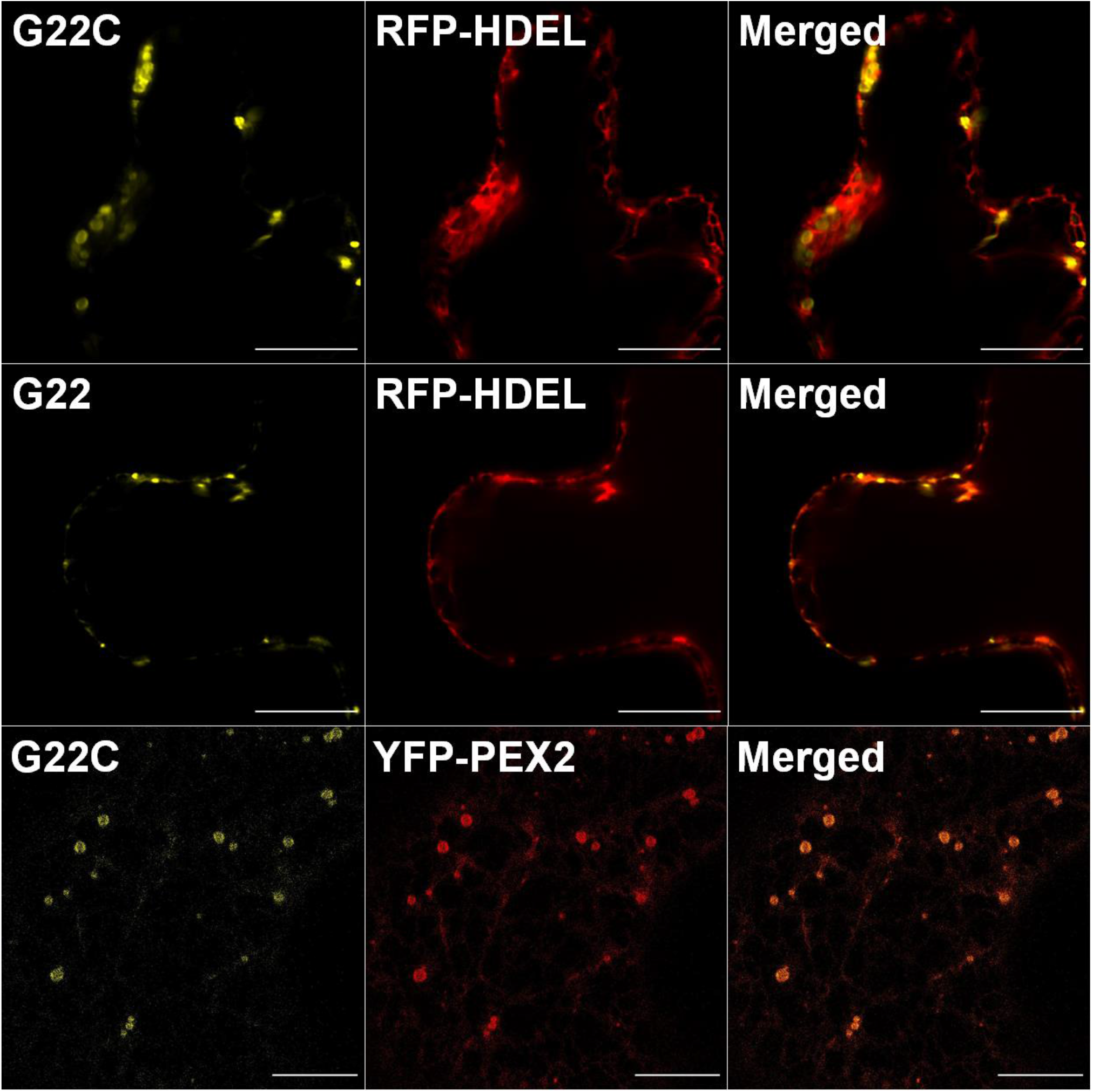
Co-expression of fluorescent markers with G22C and G22. RFP-HDEL shows no co-localisation with G22 or G22C signal. Peroxisomal marker YFP-PEX2 co-localises with G22C, suggesting a mis-localisation of the G22C proteins to the peroxisomes, instead of forming the large ER-derived compartment structures. Images were captured 7 days after agroinfiltration of mature *N. tabacum* leaves. Scale bars: 10 μm.

**Supplementary Figure S6:**
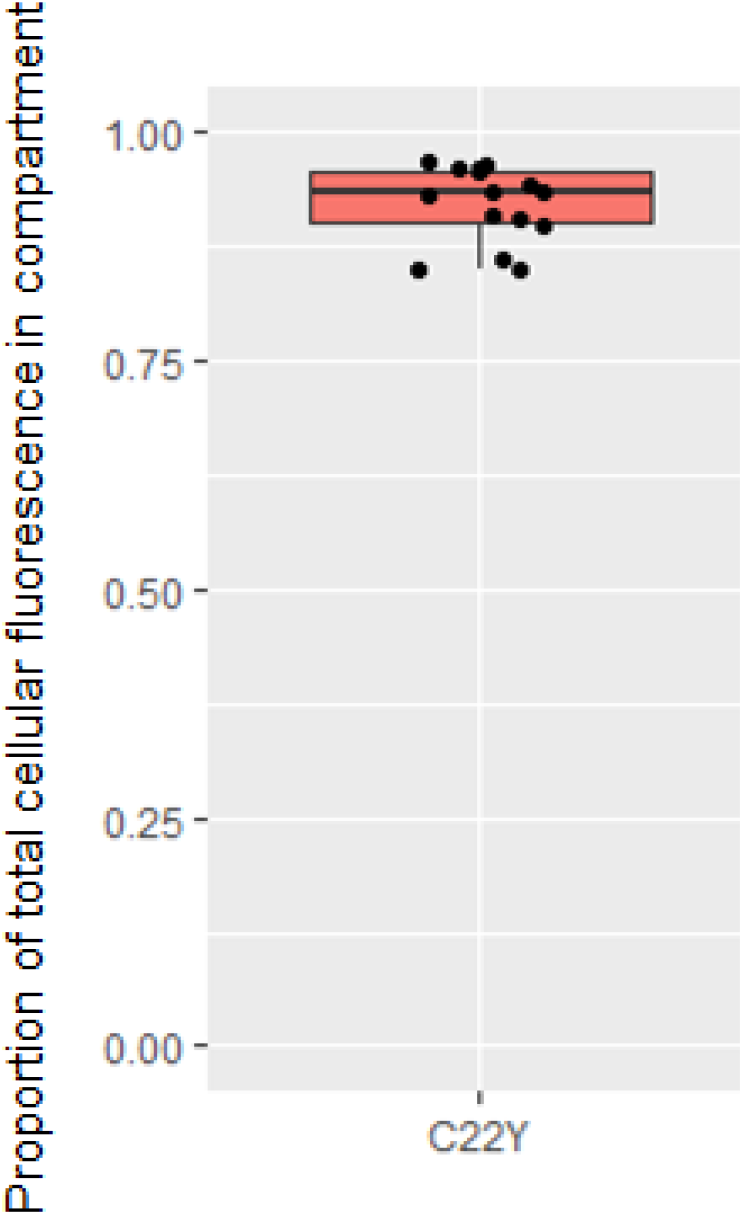
Proportion of total cellular fluorescence present in the C22Y compartment. An average of 92.7% of the total cellular YFP fluorescence is localised to the compartment, suggesting that almost all of the C22Y scaffold molecules are present in the compartment. n = 15

**Supplementary Figure S7:**
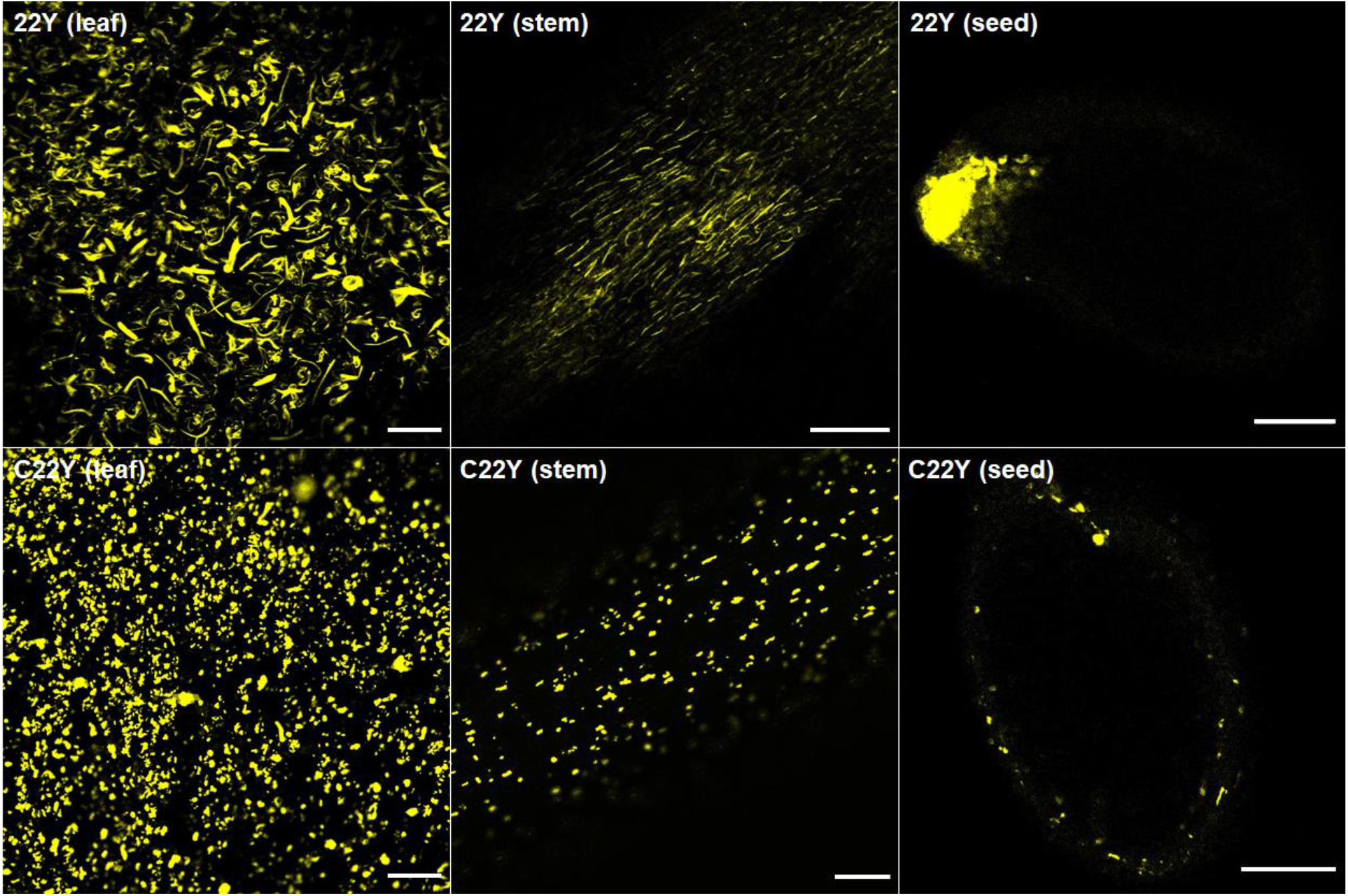
Confocal microscopy images of 6 weeks old stably transformed T2 A. thaliana lines. YFP fluorescence is consistently present in compartments in all cells for both compartment types. Seeds from both lines were also imaged, with 22Y showing faint fluorescence in the seed coat and most of the signal concentrated in the micropylar endosperm, while C22Y seeds showed several smaller compartment-like structures in the seed coat. Scale bars: 100 μm.

**Supplementary Video SV1:**
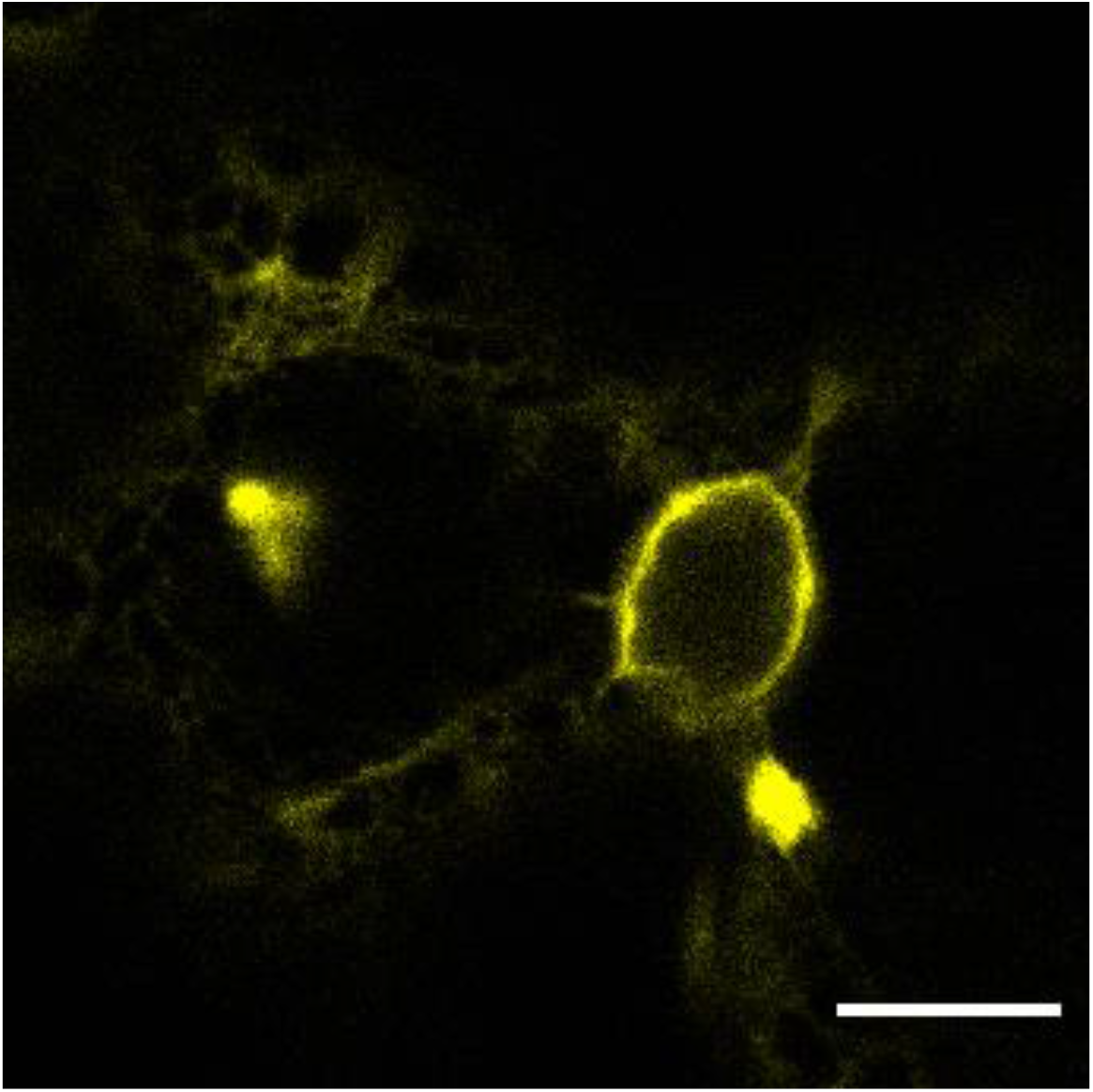
ER rearrangements in a cell with a compartment. Oversaturated YFP signal of C22Y compartments were captured to highlight the normal mobility of the ER network. Video was captured 3 days after agroinfiltration of mature *N. tabacum* leaves. Scale bar: 10 μm.

**Supplementary Table S1:**
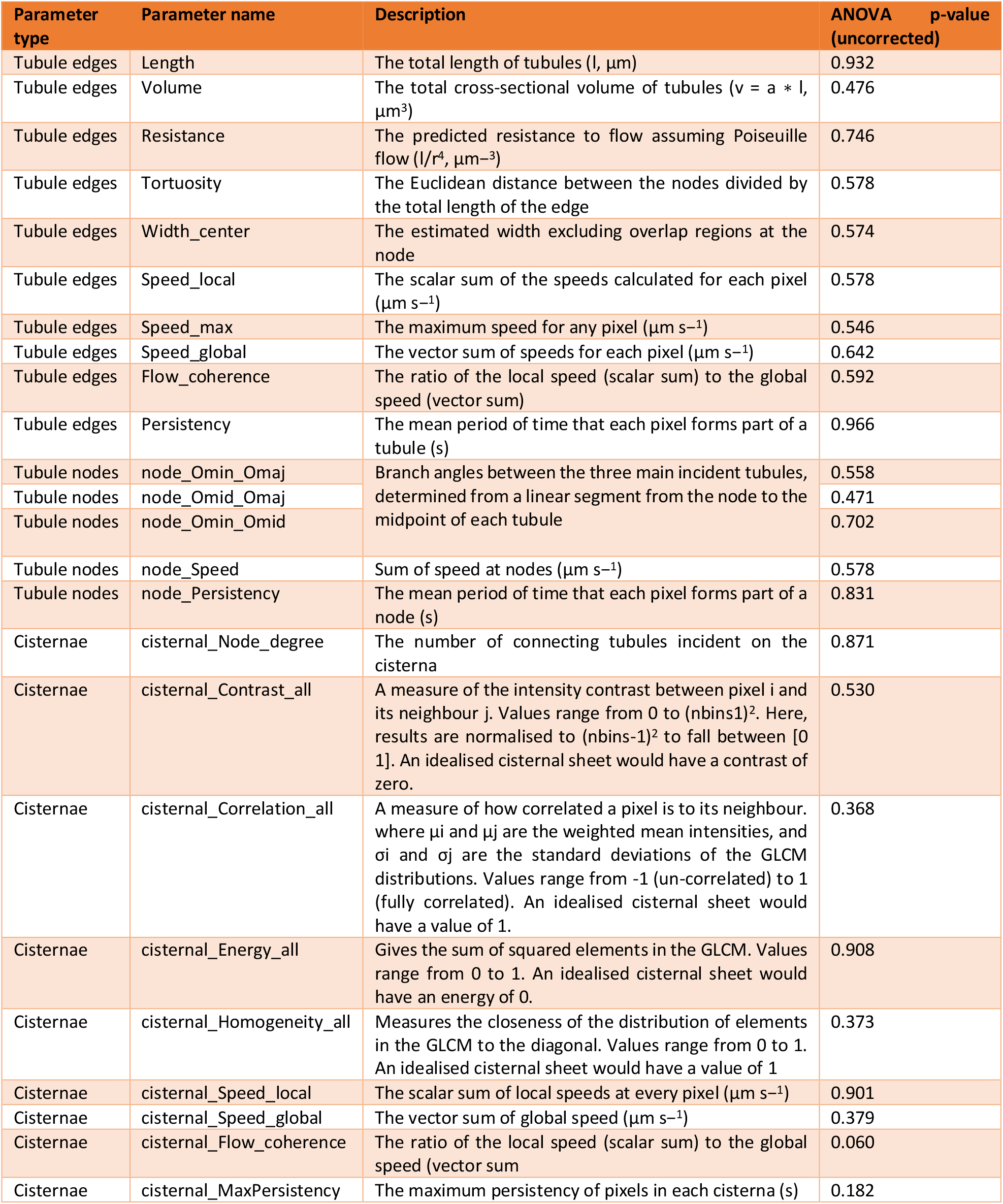
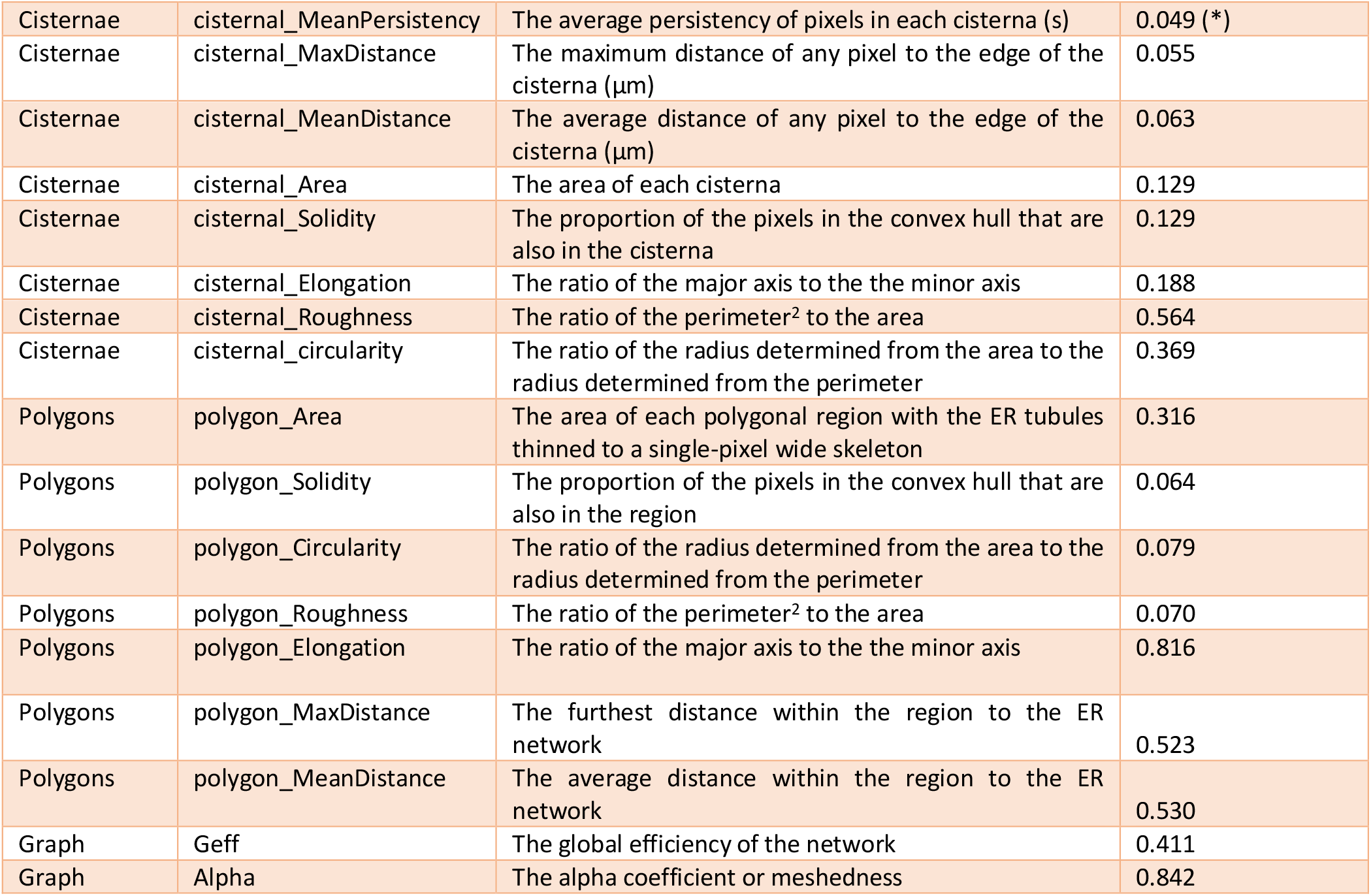
Parameters used in the AnalyzER software package to investigate ER network dynamics. Descriptions are from the AnalyzER manual (version 1.1). Individual parameters were tested using planned comparisons in ANOVA and one of these was (marked with asterisks) shown to have a statistically significant difference between at least one of the tested compartment types and the WT controls. However, when the ANOVA p-values were corrected for multiple testing, none of the parameters remained significantly different.

**Supplementary Table 2.1:**
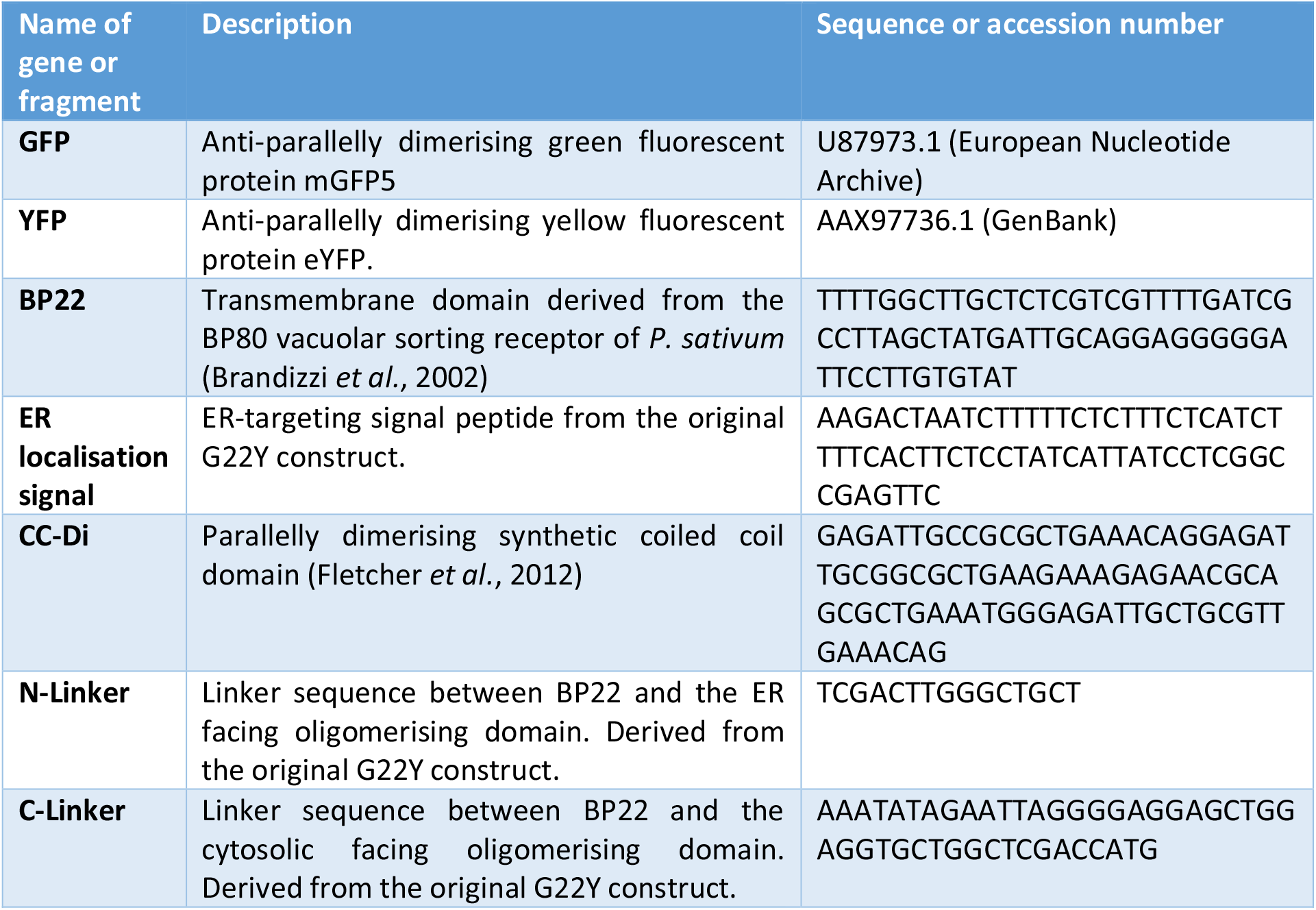
Gene parts used for the design of the genetic constructs.

**Supplementary Table 3:**
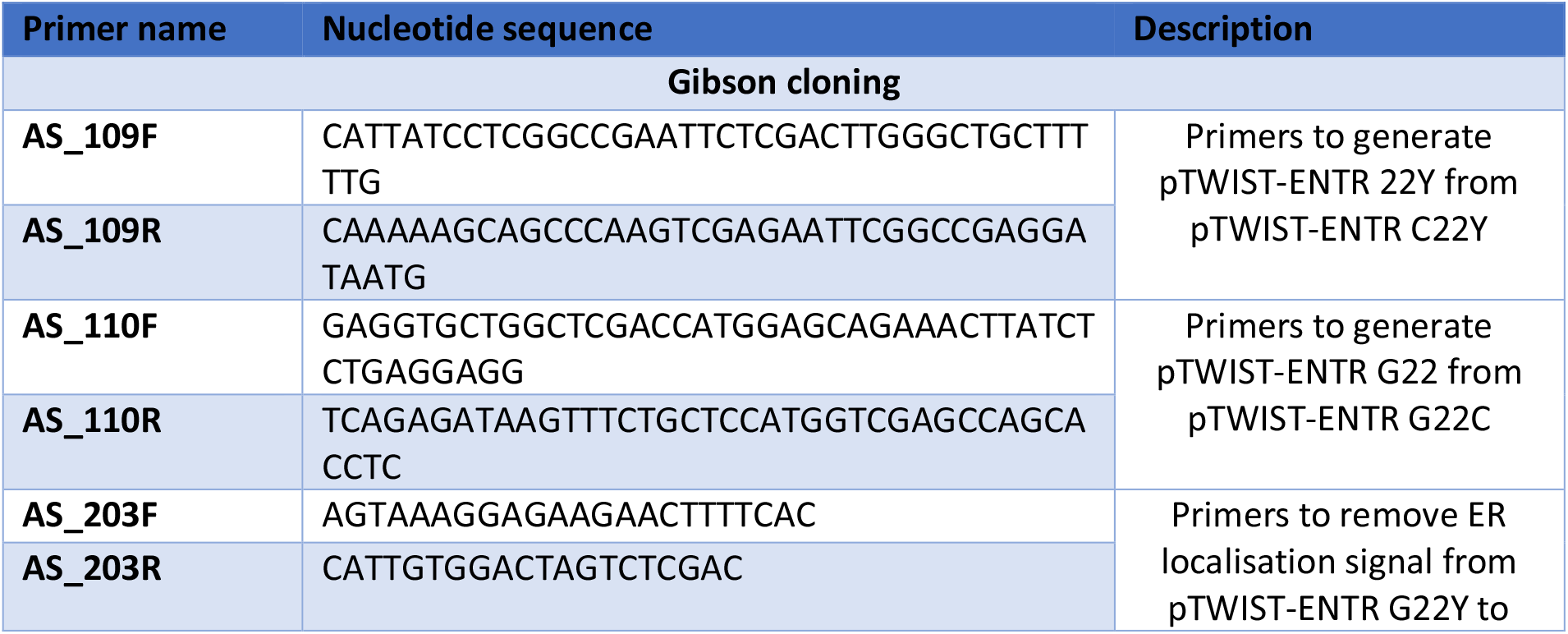
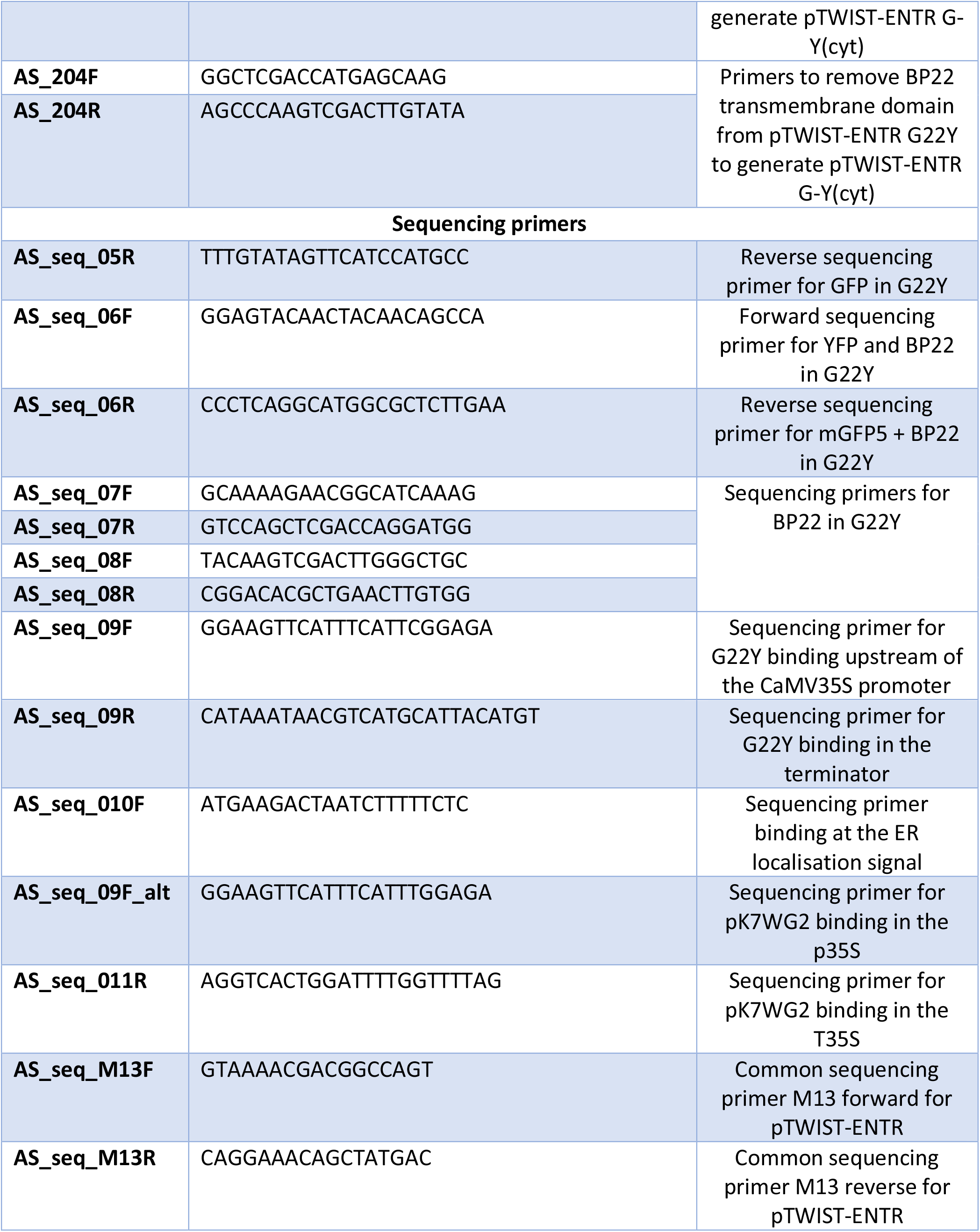
PCR and sequencing primers used to generate and confirm constructs.

